# Concurrent genetic and non-genetic resistance mechanisms to KRAS inhibition in CRC

**DOI:** 10.1101/2025.08.05.668666

**Authors:** Salvador Alonso, Kevan Chu, Marie J Parsons, Elizabeth Granowsky, Himari Gunasinghe, Jinru Shia, Rona Yaeger, Lukas E. Dow

## Abstract

KRAS is mutationally activated in 45-50% of colorectal cancer (CRC) cases, and while KRAS-targeted therapies have shown some clinical promise, upfront and acquired resistance limit their efficacy. To explore the acute response and mechanisms underlying KRAS inhibitor resistance, we used targeted exome sequencing and single-cell spatial transcriptomics to analyze patient-matched pre-treatment, on-treatment, and progression biopsies from patients treated with combined KRAS^G12C^ and EGFR inhibition. Acquired genetic events were identified in most patients at progression but were often subclonal and coexisted with transcriptional adaptive states. Mesenchymal, YAP, and fetal-like transcriptional signatures predominated in resistant tumors, while tumor cell-intrinsic inflammatory programs were induced in the early treatment phase. Single-cell spatial analysis revealed significant intratumoral heterogeneity, with diverse adaptive states predominating in different zones of individual tumors. Using human and murine organoid models, we show that these drug-induced inflammatory programs are cancer-cell autonomous and precede the emergence of regenerative fetal-like programs associated with drug resistance. We uncover TBK1 as a promising target to abrogate the early inflammatory adaptive phase and enhance responses to KRAS inhibition.

## INTRODUCTION

Colorectal cancer (CRC) is the second leading cause of cancer-related mortality worldwide, accounting for nearly 1 million deaths annually^1,2^. KRAS is the most commonly mutated oncogene in CRC, observed in up to 50% of cases^3^. The past decade has seen the emergence of multiple selective and non-selective small-molecule inhibitors targeting RAS proteins^4,5^. The first-in-class KRAS inhibitors, adagrasib and sotorasib, have demonstrated clinical benefit in patients with various cancer types harboring KRASG12C mutations^6-9^. In addition to these G12C-selective small molecules, there is now a range of additional compounds in pre-clinical and clinical development, including G12D-selective and pan-RAS inhibitors with the potential to target the majority of KRAS variants^10-13^. Thus, over the next decade, it is likely that millions of cancer patients will be treated with KRAS inhibitors.

The pivotal KRYSTAL-1 and CodeBreak 300 trials, which led to the approval of adagrasib and sotorasib, respectively, showed that only a fraction of patients with KRASG12C CRC respond to treatment^6,7^. Moreover, objective responses are often short-lived, with a median progression-free survival of less than seven months^6,7^. Analysis of post-progression cfDNA and tumor samples show that secondary RAS pathway mutations frequently emerge during treatment^14-17^. However, a subset of resistant tumors lack a clear genetic mechanism of escape. Furthermore, even when secondary mutations do emerge, they are often present at low allelic frequency and, at times, disappear under continued therapeutic pressure, indicating low clonal fitness^14-16^. Thus, while subclonal events may enable a subset of cells to escape treatment, they are unlikely to drive the bulk of resistant outgrowth.

Understanding the mechanisms underlying both primary and acquired resistance to KRAS-targeted agents is crucial for developing combination therapies that achieve deeper and more durable responses. To this end, we analyzed patient-matched clinical samples from KRAS-inhibitor trials using MSK-IMPACT targeted gene sequencing and single-cell spatial transcriptomics. Additionally, we developed resistant organoid models to functionally interrogate tumor adaptations to KRAS inhibition. Our findings reveal that both genetic and non-genetic resistance mechanisms coexist within individual patients, exhibiting significant intratumoral heterogeneity, with diverse adaptive transcriptional programs predominating in different tumor zones. Furthermore, we show that the initial acute phase of KRAS inhibition is characterized by cell-intrinsic inflammatory signaling and identify a therapeutic strategy combining KRAS plus TBK1 inhibition to mitigate these early adaptations and enhance treatment responses.

## RESULTS

### Genetic and non-genetic adaptation to KRAS inhibition in CRC

To identify genetic and non-genetic responses to KRAS inhibition, we prospectively collected 11 matched pre-treatment and on-treatment (day 7 to day 21) biopsies, as well as 8 post-progression biopsies from CRC patients treated with combined KRAS^G12C^ plus EGFR inhibition (Figure 1a, 1b). Patients received treatment in one of two clinical trials combining sotorasib plus panitumumab, or testing adagrasib with or without cetuximab^6,18^; all patients had previously received fluoropyrimidine-based chemotherapy, and all patients except one received combined KRAS^G12C^ and EGFR inhibition. Among these patients, best response consisted of stable disease in seven patients and partial responses in five patients (Figure 1a). The mean duration of treatment was 9.4 months. These outcomes closely reflect the larger trial populations^6,7^. Tumor characteristics, treatment response, and exome sequencing data (baseline and post-progression) are summarized in Figure 1a.

**Figure 1.**
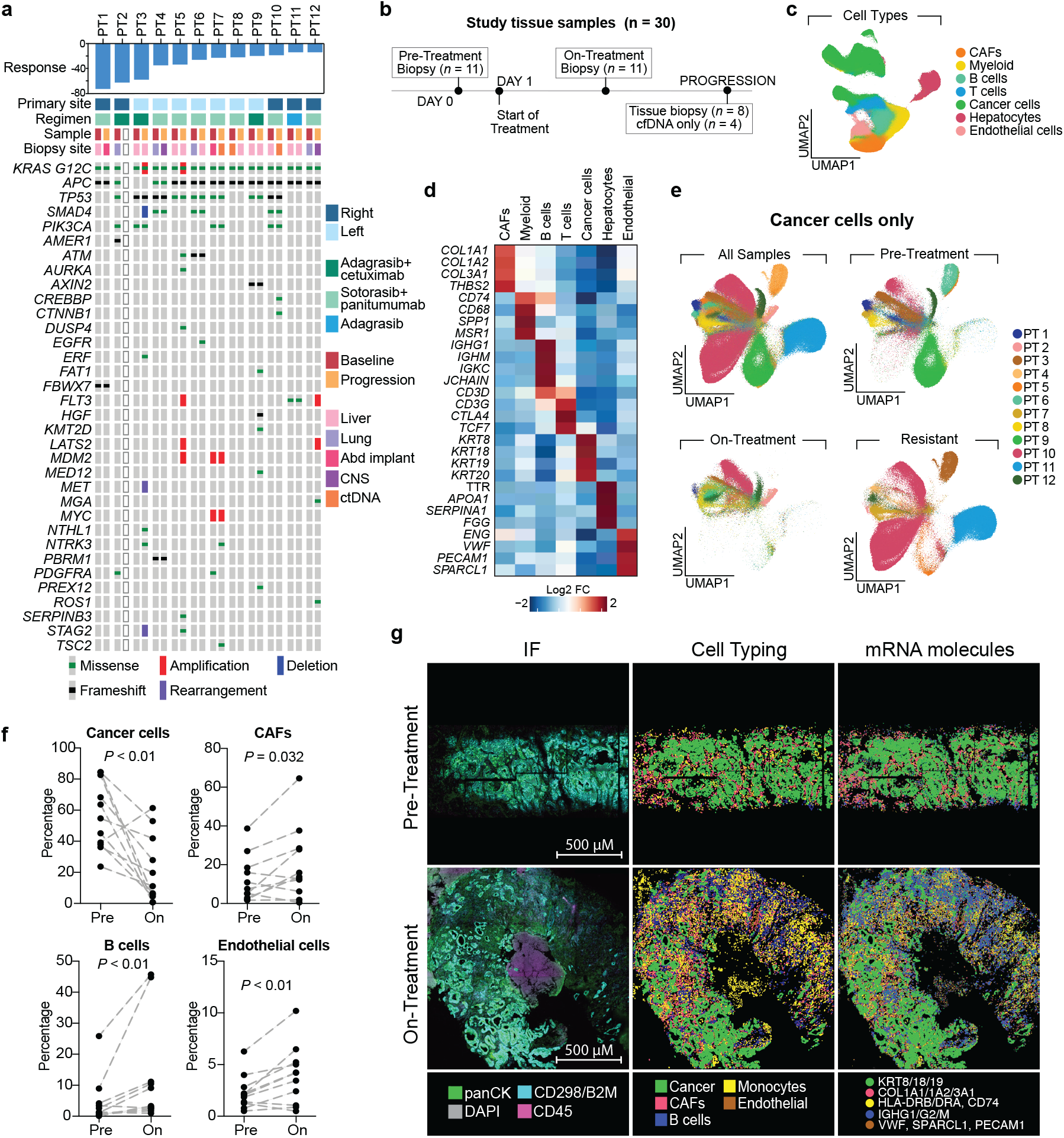
Remodeling of the tumor microenvironment in response to combined KRAS plus EGFR inhibition. **a.** Percentage change in tumor size, clinical characteristics, and oncoprint (pretreatment and post-progression) of patients with CRC treated with combined KRAS plus EGFR inhibition. **b**. Study workflow. In total, 30 pre-treatment, on-treatment, and resistant tissue biopsies were analyzed by single-cell spatial transcriptomics (Cosmx). **c**. Uniform manifold approximation and projection (UMAP) of patient samples colored by cell type. **d**. Heatmap showing normalized expression of selected cell typing markers (rows) across tumor compartments. Cancer cells were subset based on KRT8, KRT19, KRT19, and KRT20 expression. **e**. UMAP analysis of cancer cells grouped by treatment status and colored by patient ID. **f**. Percentage of cell subsets in the pre-treatment (Pre) and on-treatment (On) tissue biopsies. Connected dots represent matched patient samples. P values were calculated using a two-tailed Wilcoxon rank sum test. **g**. Representative pre-treatment and on-treatment patient-matched samples. Left: Immunofluorescence images. Middle: Cell typing inferred from transcriptomic data. Dots represent individual cells. Right: Spatial localization of cell typing transcripts. Dots represent individual transcripts.

Targeted exome sequencing using MSK-IMPACT showed acquired genetic events in seven out of the eleven patients for whom pre-treatment and post-progression samples were available (including solid tumor or liquid biopsies) (Figure 1a and S1). Six patients had one or more acquired mutations in known regulators of RAS signaling, including KRAS^G12C^ and FLT3 amplifications, and missense mutations in EGFR, FLT3, DUSP4, and ROS1. The majority of putative resistance drivers were subclonal; of the 17 de novo mutations detected in progression biopsies, only seven were clonal (defined as a variant allele frequency > 50% relative to preexisting clonal mutations) (Figure S1). There were no identifiable acquired genetic events in the remaining four patients. Thus, similar to what was previously reported in CRC and other cancer types following progression on KRAS inhibitors, escape mutations were not universally identified and were often subclonal^14-16^. To identify non-genetic tumor adaptations, we performed spatial transcriptomics of matched pre-treatment, on-treatment, and progression biopsies (Figure 1b). We employed a cyclic fluorescent in situ hybridization platform (CosMx) that quantifies the relative abundance of 1000 transcripts at single-cell resolution directly on formalin-fixed paraffin-embedded (FFPE) tissue^19^. A total of 933,903 cells passed quality control across all patient samples, including 409,858 cancer cells. Unsupervised clustering of single-cell profiles identified 14 cell subsets, which were annotated by known gene expression markers (Figure 1c, 1d, and S2a). Cancer cells were annotated based on their high expression of keratin genes (KRT8, KRT18, and KRT19) (Figure 1d). Malignant cells showed significant heterogeneity, and generally clustered by patient and treatment status (Figure 1e). Pre-treatment and resistant samples were transcriptionally more diverse compared to on-treatment cancer cells, which generally clustered together (Figure 1e). As expected, analysis of cell type abundance showed a significant decrease in the number of malignant cells early on-treatment relative to the pre-treatment time points (Figure 1f and 1g). There was also a higher proportion of cancer-associated fibroblasts (CAFs), B cells, and endothelial cells in the on-treatment relative to pre-treatment biopsies (Figure 1f, 1g, and S2b).

To identify adaptive responses to RAS inhibition specifically in tumor cells, we examined differentially expressed genes and programs in the malignant compartment of patient-matched biopsies. Over-representation analysis (ORA) across Hallmark, KEGG, and Reactome collections showed activation of inflammatory programs in on-treatment and progression samples (Figure 2a). Similarly, Gene Set Variation Analysis (GSVA) on individual samples using hallmark and intestinal cell type-specific gene sets showed an increase in inflammatory pathways and intestinal stem cell signatures (Figure 2b). Annotation of individual genes that were significantly differentially expressed (Log2 fold-change > 0.3, adjusted p <0.05) in at least two thirds of the samples again showed a marked enrichment of interferon and cytokine signaling in on-treatment samples (Figure 2c). In contrast, therapy-resistant (progression) samples showed an enrichment of epithelial-to-mesenchymal transition (EMT), YAP, and fetal-like programs (Figure 2b, 2c, and S3-4) that we and others have previously linked to CRC metastasis and WNT-targeted therapy resistance^20-30^. There was also a marked downregulation of epithelial and intestinal differentiation markers, and corresponding increase in stem cell markers under KRAS inhibitor treatment, but in most cases, these signatures returned to baseline in progression samples (Figure 2b). As expected, the expression of KRAS targets was lower on-treatment compared to baseline and subsequently increased at progression in a subset of patients (Figure 2b, 2c, and S4), likely due to the withdrawal from KRAS/EGFR therapy.

**Figure 2.**
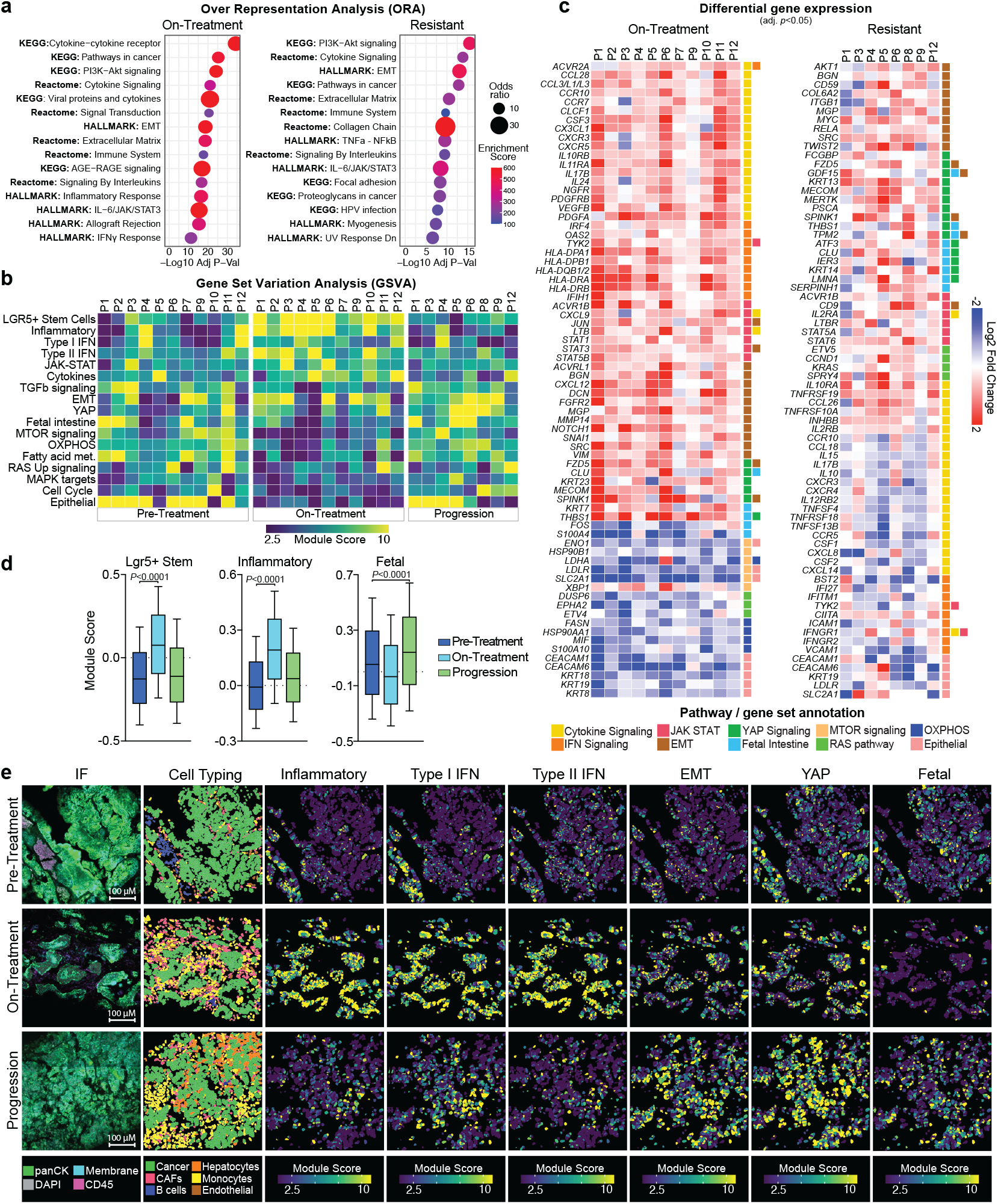
Transcriptomic adaptations to KRAS inhibition in the cancer cell compartment. **a.** ORA bubble plot showing the five most significantly enriched pathways from the Hallmark, Reactome, and KEGG gene set collections in the on-treatment versus pre-treatment, and resistant versus pre-treatment patient samples. Differentially expressed genes were identified using a Wilcoxon rank sum test. **b**. Heatmap displaying module score quantiles of Hallmark and intestinal cell type-specific gene sets across individual patient samples. **c**. Cancer cell differential gene expression in the on-treatment and resistant (versus pre-treatment) patient-matched samples. Genes for which Log2FC > 0.3 or < -0.3 in at least two thirds of samples are displayed. Colored squares represent the signaling pathways associated with each gene. **d**. Module scores of selected gene sets in cancer cells according to treatment timepoint. Error bars show the 10-90 percentiles. P values were calculated using a two-tailed Wilcoxon rank sum test. **e**. Representative case (patient 6) showing immunofluorescence and spatial images. The gradient scale shows the module score quantiles of resistance programs in cancer cells (non-malignant cells were removed from the analysis).

A notable outlier in our data was patient 4. In contrast to other cases, the pretreatment biopsy showed robust expression of type I and type II IFN programs, which were downregulated following initiation of sotorasib plus panitumumab (Figure S4). Importantly, before the on-treatment biopsy, the patient started treatment for a flare of rheumatoid arthritis with the anti-inflammatory drug hydroxychloroquine and also received rituximab during the study treatment. The patient had an objective response and derived meaningful clinical benefit, remaining on treatment for 23.5 months — the longest duration observed in our study cohort (median time on treatment: 9.4 months).

### Coexisting genetic and non-genetic resistance to KRAS inhibition

The presence of both genetic and non-genetic transcriptional changes in progression samples raised the question of whether these cellular changes were mutually exclusive resistance mechanisms. In fact, transcriptional reprogramming was identified in each of the eight post-progression biopsies analyzed, including those with multiple putative genetic resistance alterations (Figure 3a, 3b, and S4). Post-progression tumor exome sequencing of patient 5, for instance, showed amplification of KRAS^G12C^ (5.9-fold copy number gain), FLT3 and MDM2, in addition to acquired missense mutations in DUSP4, ATM, AURKA, SERPINB3, and STAG3 (Figure 3a). In addition to the expected upregulation of KRAS transcripts and downstream reactivation of MAPK signaling, likely driven by the KRAS^G12C^ amplification and other RAS pathway mutations, transcriptomic analysis revealed strong enrichment of type I IFN, YAP and fetal intestinal signatures at progression (Figure 3b), revealing that even in cases with classic genetic resistance profiles, transcriptional and lineage adaptations occur.

**Figure 3.**
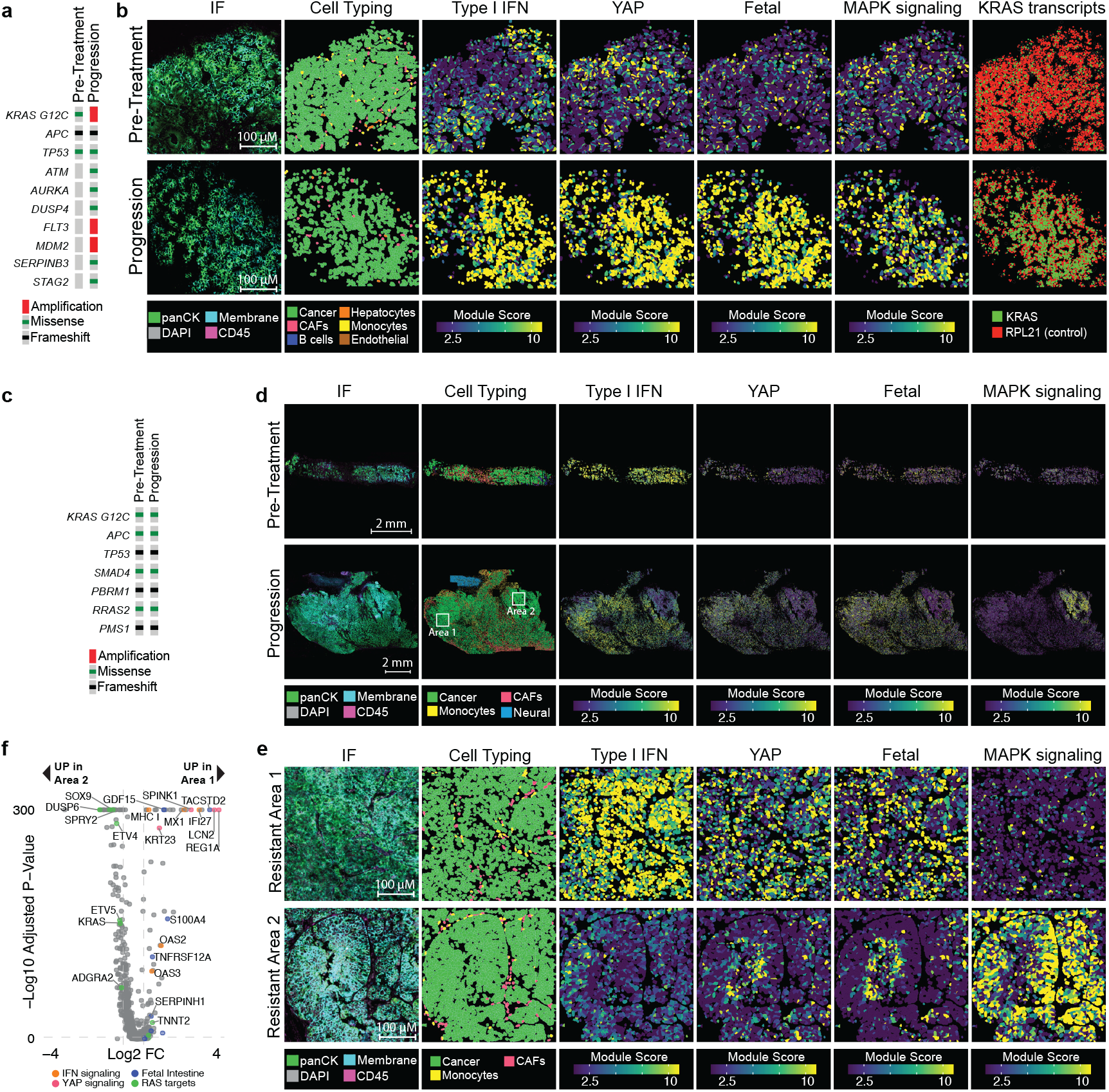
Co-existing genetic and non-genetic resistance mechanisms to KRAS inhibition. **a.** Oncoprint of the pre-treatment and post-progression biopsies from patient 5. **b**. Immunofluorescence and spatial images of the pre-treatment and resistant biopsies from patient 5. The gradient scale shows module score quantiles of selected gene sets in cancer cells. Panels in the far-right show transcripts for RPL21 (endogenous control) and KRAS (each dot represents an individual transcript). **c**. Oncoprint of the pre-treatment and post-progression biopsies from patient 4. **d**. Immunofluorescence and spatial images of patient 4 observed at low magnification. Two areas in the post-progression biopsy with seemingly divergent resistance mechanisms are highlighted. **e**. Area 1 and Area 2 are displayed at higher magnification. The gradient scale shows module score quantiles of selected gene sets in cancer cells. **f**. Volcano plot showing differentially expressed genes in Area 1 versus Area 2. Differentially expressed genes corresponding to IFN, YAP, fetal, and RAS programs are highlighted. P values were calculated using a two-tailed Wilcoxon rank sum test with Benjamini-Hochberg correction for multiple comparisons.

In some cases, the MSK-IMPACT sequencing and transcriptomic data revealed co-existing genetic and non-genetic resistance mechanisms, but the spatial analysis revealed significant intratumoral heterogeneity, with enrichment of distinct adaptive programs in different zones of individual tumors (Figure 3d, 3f, and S4-5). Patient 4, for instance, progressed after 23.5 months of treatment with sotorasib plus panitumumab, and MSK-IMPACT sequencing showed no genetic events predicted to activate MAPK signaling (Figure 3c). Spatial transcriptomics revealed two distinctive areas with seemingly mutually exclusive adaptive resistance mechanisms (Figure 3d-f). Area 1 showed strong enrichment of type I IFN, YAP, and fetal intestinal programs but low MAPK signaling. In contrast, Area 2 showed robust MAPK reactivation but suppressed type I IFN, YAP, and fetal markers (Figure 3d-3f). These observations suggest that independent resistant clones may evolve within the same refractory tumor and may explain the frequent emergence of multiple subclonal genetic events at progression captured through cell-free DNA profiling^14-16^.

### Cancer cell-autonomous activation of inflammatory programs following KRAS inhibition

We next asked whether transcriptional reprogramming is dependent on the tumor microenvironment. To investigate potential cell-cell communications, we performed ligand-receptor interaction analysis using CellChat^31^, which infers intercellular signaling by estimating the probability of interaction based on the expression levels of ligands in sender cells and corresponding receptors in receiver cells. Despite transcriptional upregulation of various inflammatory mediators in the non-malignant compartment, there was minimal predicted ligand-receptor signaling directed toward cancer cells. Only two weak but statistically significant interactions were identified: SPP1-CD44 and LGALS9-CD44, both involving monocyte-derived ligands (SPP1 and LGALS9) engaging the receptor CD44 on malignant cells (Figure 4a and S6a). These data suggest that while inflammatory genes are transcriptionally induced in the microenvironment, direct ligand-receptor signaling from immune to cancer cells may be limited. However, we cannot exclude that microenvironmental ligands not included in the 1000-gene panel activate inflammatory programs in cancer cells.

**Figure 4.**
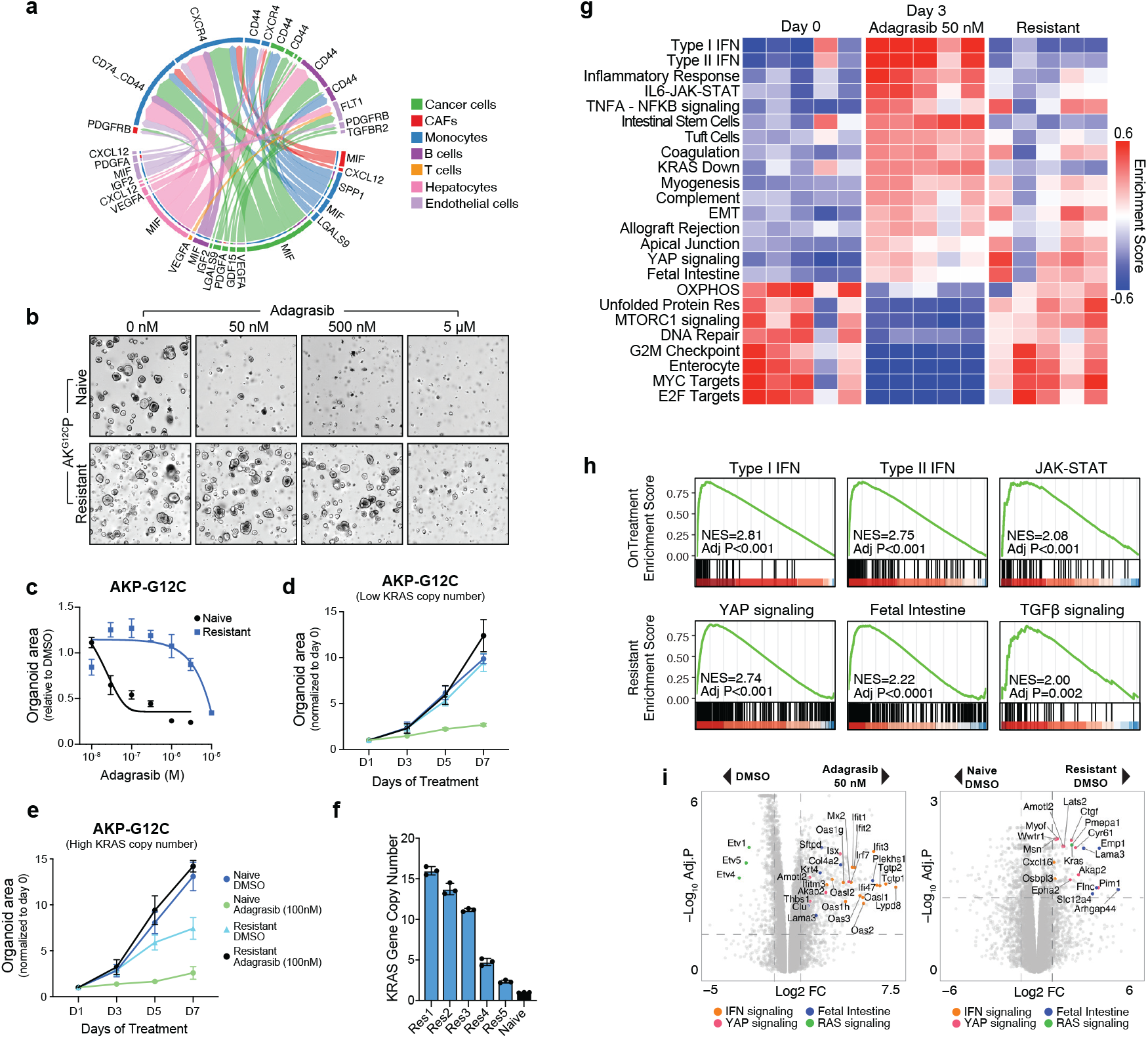
Transcriptomic adaptation to KRAS inhibition is cancer-cell autonomous. **a.** CellChat chord diagram displaying ligand-receptor interactions between cell compartments in patient biopsies. Arrows indicate the directionality of the interaction. Wider connections correspond to stronger signaling. Only interactions with P value <0.05 (permutation test with Benjamini-Hochberg correction) are displayed. **b**. Representative brightfield images of treatment-naïve and KRAS-inhibitor resistant AKP-G12C organoids treated with increasing concentrations of adagrasib. Images were obtained on day 3 of treatment. **c**. Growth of treatment-naïve and resistant AKP-G12C organoids treated with increasing concentrations of adagrasib. Organoid growth was normalized to untreated conditions. **d**. KRAS copy number of treatment-naïve and KRAS-inhibitor resistant AKP-G12C organoids. **e**. Growth of treatment-naïve and resistant AKP-G12C organoids (low KRAS copy number) treated with the EC80 of adagrasib (100 nM). **f**. Growth of treatment-naïve and KRAS-inhibitor-addicted (high KRAS copy number) AKP-G12C organoids treated with the EC80 of adagrasib (100 nM). **g**. Heatmap showing GSEA enrichment scores for significantly enriched or depleted Hallmark gene sets in AKP-G12C organoids following treatment with DMSO, adagrasib 50 nM (EC50 for 72 hours), or at resistance. **h**. GSEA plots showing significantly enriched Hallmark gene sets following treatment with adagrasib (50 nM for 72 hours) (top row), or at resistance (bottom row), relative to DMSO-treated AKP-G12C organoids. P values were calculated using a permutation test with Benjamini-Hochberg correction. **i**. Volcano plot showing differentially expressed genes in the on-treatment (adagrasib 50 nM for 72 hours) versus DMSO-treated, and resistant versus naïve AKP-G12C organoids. Select differentially expressed genes corresponding to IFN, YAP, fetal, and RAS programs are highlighted. P values were calculated using a two-tailed Wilcoxon rank sum test with Benjamini-Hochberg correction. For organoid experiments (c, d, f), P values were calculated using a two-tailed t test (n = 4 independent replicates). Error bars represent the standard error of the mean. ^**^, P < 0.01.

To model adaptive responses to KRAS inhibition in the absence of a complete tumor microenvironment, we generated C57BL/6 colon organoids carrying an *Apc* truncation (APC^Q1405X^), an endogenous KRAS^G12C^ mutation, and a *Trp53* truncation (p53^Q97X^) (hereafter AKP-G12C). AKP-G12C organoids showed rapid growth arrest following KRAS inhibition (adagrasib 50 nM); however, a subset survived and gave rise to highly resistant organoids after 10 to 12 weeks of treatment (Figure 4b). Resistant organoids showed an EC50 shift of 50 to 100-fold compared to treatment-naïve organoids (Figure 4c). Adagrasib-resistant organoids were cross-resistant to the KRAS^G12C^ inhibitor sotorasib, and the recently described multi-selective RAS(ON) inhibitor RMC-7977 (Figure S6b and S6c).

To characterize the response to KRAS inhibition, we performed RNA sequencing of AKP-G12C organoids prior to, during treatment (adagrasib 50 nM for 72 hours), and at progression, mirroring the timepoints of patient biopsies. In total, we generated five independent adagrasib-resistant AKP-G12C lines. Four resistant lines showed strong enrichment of KRAS transcripts and its downstream targets (Figure S6d). Interestingly, these resistant lines were addicted to KRAS inhibition, as drug withdrawal led to rapid growth arrest (Figure 4d and 4e). These findings were reminiscent of the senescent phenotype observed upon drug withdrawal in KRASG12C-amplified CRC cells^16^. Indeed, copy number analysis confirmed KRAS amplifications in each of the KRAS-inhibitor-addicted lines (mean copy number 6, range 4.8-16 copies) (Figure 4f).

Transcriptome analysis revealed enrichment of inflammatory, EMT, fetal intestine, and YAP signatures in resistant organoids, independent of KRAS copy number alterations, demonstrating coexistence of genetic and non-genetic escape drivers, as observed in the clinical samples (Figure 4g-i). Inflammatory programs, including type I and type II IFN, JAK-STAT, and TNFα-NFκB were strongly enriched on-treatment, and remained modestly elevated in resistant organoids, albeit at lower levels (Figure 4g-i). These findings also mirrored our data in patient biopsies, and indicate that inflammatory programs, generally associated with refractory tumors, are engaged early following KRAS inhibition^20-25^. Further, the organoid data demonstrate that reprogramming is at least partially cancer cell-autonomous and not strictly dependent on non-malignant cells within the tumor microenvironment.

### TBK1 inhibition blocks early inflammatory reprogramming and synergizes with KRAS inhibition

Our group and others have shown that inflammatory programs are engaged during the initial response of cancer cells to targeted and cytotoxic therapies^20-24,32^. However, whether these programs are involved in the adaptive process or are passenger signs of cellular stress remains unknown. To test whether blockage of inflammatory signaling would abrogate initial adaptive responses and synergize with KRAS inhibition, we conducted a small, focused drug screen in KRAS^G12C^ patient-derived organoids (PDOs) combining adagrasib with kinase inhibitors targeting inflammatory pathways. We evaluated two concentrations (0.5 and 1 µM) of small molecules targeting eight kinases: PI3K, FAK, KIT, JNK, IKK, TGF?RI, JAK, and TBK1 (Figure 5a and 5b). Of these, only the TBK1 inhibitor TBK1/IKKX-IN-5 delayed outgrowth in the presence of adagrasib (Figure 5a and S7a); TBK1 inhibition alone had no significant effect on organoid growth. Similar results were observed with the FDA-approved dual TBK1/JAK inhibitor momelotinib (Figure 5b). TBK1 and the related kinase IKKX activate the transcription factor IRF3, a key mediator of the innate antiviral immune response^21,33^. Analysis of patient-matched samples revealed that IRF3 target genes included in the CosMx panel were upregulated following combined KRAS and EGFR inhibition (Figure 5c and S7b). Similarly, differential gene expression and gene set analysis of AKP-G12C organoids treated with adagrasib showed robust enrichment of IRF3 targets (Figure 5d and S7c), supporting the notion that TBK1–IRF3 signaling is triggered in response to KRAS inhibition. To ensure the combined effect of adagrasib and TBK1 inhibition was not specific to a single PDO line, we treated a second KRAS^G12C^ PDO, as well as AKP murine organoids carrying either KRAS^G12C^ or KRAS^G12D^ mutations (Figure 5e, S7d and S7e). In each case, TBK1 blockade had minimal impact as a monotherapy, but stunted organoid growth in combination with KRAS inhibition (Figure 5e, S7d, and S7e). Both TBK1/IKKX-IN-5 and momelotinib inhibit related kinases in addition to TBK1, however silencing of only TBK1 using two independent doxycycline-inducible shRNAs mimicked the effect of drug treatment (Figure 5f and 5g), implying that TBK1 inhibition is sufficient to enhance the activity of adagrasib treatment on drug-naïve organoids. Together, these data highlight that the acute induction of inflammatory programs following KRAS inhibition, at least in part through TBK1, may play a role in the emergence of drug-resistant tumor outgrowth.

**Figure 5.**
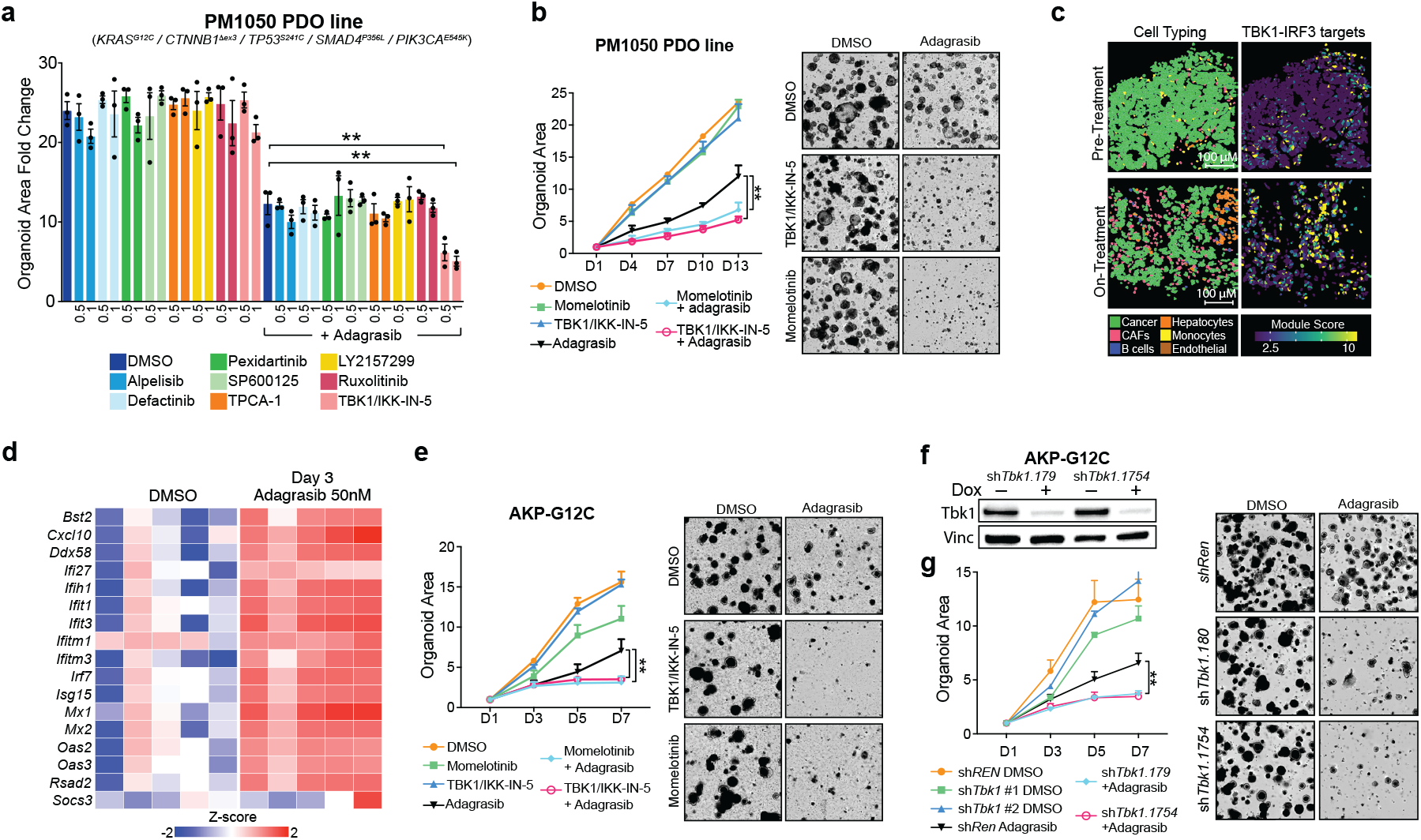
Targeting early inflammatory reprogramming via TBK1 blockage synergizes with KRAS inhibition. **a.** Bar plot showing focused drug screen results of PM1050 PDOs treated with adagrasib alone (50 nM), kinase inhibitors targeting inflammatory programs, or the combination. Two concentrations of each kinase inhibitor were tested. Bar values represent the organoid area fold change during the 13-day experiment. The x-axis shows the concentration of the kinase inhibitor (µM). **b**. Growth curve and brightfield images of PM1050 organoids treated with KRASi (adagrasib 50 nM), TBK1i (TBK1/IKK -IN-5 1 µM, momelotinib 1 µM), or the combination. **c**. Representative case (patient 5) showing images of the pre-treatment and on-treatment samples. The gradient scale shows the module score quantiles of TBK1-IRF3 targets (only cancer cells are displayed). **d**. Differential gene expression analysis (Z-score) of TBK1-IRF3 targets in AKP-G12C organoids following treatment with DMSO, or adagrasib 50 nM (EC50 for 72 hours). **e**. Growth curve and brightfield images of AKP-G12C organoids treated with KRASi (adagrasib 50 nM), TBK1i (TBK1/IKK -IN-5 0.5 µM, momelotinib 1 µM), or the combination. **f**. Western blot of AKP-G12C organoids transduced with dox-inducible shRNAs targeting Tbk1 (shTbk1). **g**. Growth curve and brightfield images of shTbk1 and shRen (control) AKP-G12C organoids treated with adagrasib (50 nM), doxycycline (1 µg/mL), or the combination. P-values were calculated using a two-tailed t-test (n = 4 independent replicates). Error bars represent the standard error of the mean. Brightfield images show organoid density on the final day of the experiment. ^**^, P < 0.01, ^*^, P < 0.05.

## DISCUSSION

Adagrasib and sotorasib have demonstrated significant clinical benefit in treating advanced epithelial cancers, and several next-generation RAS inhibitors are now in late-stage clinical development, offering the potential to significantly improve outcomes for millions of cancer patients in the coming years. However, as with other targeted therapies, primary or acquired resistance is likely to occur in virtually all patients. Understanding the mechanisms underlying drug escape is essential for developing strategies to delay, or even prevent, treatment failure. Recent studies show that KRAS inhibitor-resistant tumors frequently acquire de novo mutations predicted to reactivate RAS signaling^14-17^. In agreement with recent studies, our analysis identified acquired RAS pathway mutations in the majority of progression samples; however, as in previous examples^14-17^, most mutations were subclonal, raising the question of whether they are bona fide drivers of resistant outgrowth.

By integrating genomic and transcriptomic data from serially collected, patient-matched biopsies and cancer models, our study reveals that acquired genetic alterations often co-exist with non-genetic resistance mechanisms. We show that resistant tumors engage YAP-associated transcriptional programs reminiscent of regenerative or fetal-like intestine, which we and others have reported to be associated with metastasis and multidrug resistance^20-30,34,35^. In some tumors, these adaptive responses are engaged in parallel to MAPK pathway reactivation, likely driven by genetic events, such as KRAS^G12C^ amplifications^16^. Notably, transcriptional programs in the post-progression biopsies were more diverse compared to the pre-treatment and on-treatment samples. Indeed, we observed significant intratumoral heterogeneity at progression with inflammatory responses, EMT, YAP signaling, fetal-like programs, and MAPK pathway reactivation predominating in different zones of individual tumors, suggesting that clones evolve independently via the engagement of diverse escape programs. In other cases, genetic and non-genetic programs co-existed within the same tumor zones and individual cells. These findings may explain the detection of multiple subclonal events in post-progression cfDNA samples, which, as opposed to tumor biopsies, capture genetic drivers across a patient’s metastatic tumors^14-17^.

The analysis of on-treatment samples revealed transcriptional rewiring as early as seven days following treatment initiation, or as early as three days in ex vivo organoid culture. Notably, on-treatment programs were significantly less diverse compared to resistant phenotypes and often involved inflammatory signatures previously shown to play a role in therapy resistance^20-25,34,35^, indicating they may serve as early mediators of tumor adaptation to KRAS inhibition. Focused drug screens targeting kinases involved in inflammatory programs showed that combined KRAS plus TBK1 inhibition restrained the growth of patient-derived and murine organoids. This is not the first indication of a role for TBK1 inhibition in KRAS mutant cancers^36,37^; However, in contrast to previous work by Barbie et al suggested a specific TBK1-dependency in KRAS mutant cells, our work implies that TBK1 is engaged following KRAS inhibition and that targeting both proteins enhances the anti-tumor efficacy. These results align with recent studies in EGFR-mutant non-small cell lung cancer, where EGFR inhibition activates an innate antiviral response through the TBK1–IRF3 axis, and combined TBK1 and EGFR inhibition yields synergistic effects in preclinical models^21^.

While our study sheds light on critical adaptive processes and the significant heterogeneity of drug resistance mechanisms within individual tumors, there are limitations. The relatively small cohort size limits our ability to correlate specific clinical or genomic profiles with transcriptional adaptations. Furthermore, due to restrictions on sample volume, the scope of transcriptional and epigenetic profiling is limited, constraining our ability to dissect rare malignant or immune populations, and the analysis of tumor-stroma interactions. Nonetheless, the reproducibility of key inflammatory signatures across samples and organoid models underscores the relevance of these adaptive mechanisms.

In all, our study reveals the complexity of the adaptive response to KRAS inhibition in CRC. The inter- and intratumoral heterogeneity in progression samples, and the coexistence of genetic and non-genetic resistance mechanisms, underscores the challenge of reversing therapeutic resistance. Targeting early inflammatory adaptations, which we show to be more conserved and generalizable within and across tumors, may offer a path for deeper and more durable responses to RAS-targeted therapies in CRC.

### Contributions

S.A., R.Y., and L.E.D. conceived the project. S.A. and K.C. performed experiments and analyzed data. M.J.P. developed the murine colon cancer organoids. E.G. and H.G. performed experiments. J.S. performed pathologic analysis of patient samples. S.A. and L.E.D. supervised experiments, performed computational work, analyzed data, and wrote the manuscript.

### Conflicts

LED is a consultant and has received research funding from Revolution Medicines, unrelated to this work. LED and Weill Cornell have licensed technologies related to KRAS to Boehringer Ingelheim, Amgen, and Revolution Medicines. RY has served as an advisor for Mirati Therapeutics, Revolution Medicine, Loxo@Lilly, Merck, and Erasca, has received a speaker’s honorarium from Zai Lab, and has received research support to her institution from Pfizer, Boehringer Ingelheim, Boundless Bio, Mirati Therapeutics, Daiichi Sankyo, Amgen, and Revolution Medicine.

## Acknowledgments

We thank the Integrated Genomics Operation (supported by grant P30 CA008748) and the Marie-Josée and Henry R. Kravis Center for Molecular Oncology at Memorial Sloan Kettering. This work was supported by the Starr Cancer Consortium (I16-0062), the Emerald Foundation, a K08 Career Development Award (1K08CA279499) and a K12 Paul Calebresi Award (K12 CA184746) from the NCI (S.A.), a Career Award for Medical Scientists from the Burroughs Wellcome Fund (Award ID 1250167) (S.A.), a Young Investigator Award from the Conquer Cancer Foundation (S.A.), and a Burroughs Wellcome Fund Physician-Scientist Institutional Award (G-1020043) (S.A.) R.Y. was supported by an NIH R21 (1R21CA292178). LED was supported by an Emerald Foundation Distinguished Investigator Award.

## METHODS

### Cloning

PCR-amplified 97mer oligos were inserted into EcoRI and XhoI-digested pCF80638 (Addgene 186711) by standard restriction cloning. Vectors and inserts were ligated using T4 DNA ligase (NEB M0202M) at 16 °C overnight and transformed into Stbl3 chemically competent cells. Oligos for shRNA cloning are listed in Supplementary Table 1. For lentiviral production, HEK293T cells were transfected with the appropriate lentiviral transfer vector along with the packaging plasmids psPax2 (Addgene, catalog no. 12260) and pMD2.G (Addgene, catalog no. 12259) using polyethylenimine. Viral supernatants were harvested at 48, 72, and 96 hours post-transfection, centrifuged at 25,000 RPM and 37°C for 2 hours, and stored at –80°C.

### Cell lines

HEK293T cells were purchased from ATCC (CRL-3216). Stocks were tested for mycoplasma routinely every six months and maintained in DMEM (Corning, 10-013-CV) containing 1% Pen/Strep (Fisher Scientific 15140163) and 10% FBS (Gibco A5670801) at 37 °C with 5% CO2.

### Murine colon cancer organoids

Mice were euthanized by inhaled carbon dioxide and sprayed down with 70% ethanol. The abdominal cavity was opened, the colon was removed and opened longitudinally with sterile scissors. Feces and excess mucus were scraped off with a slide. The colon was transferred to a 50 mL tube and washed with 20 mL of ice-cold PBS. The colon was then cut into 5–10 mm pieces, placed in 10 mL of ice-cold EDTA/PBS, and incubated on a shaker at 4°C for 60 minutes. Following incubation, the tube was shaken vigorously 10 times, and the supernatant was removed. The tissue was resuspended in 20 mL of ice-cold sterile PBS containing 200 U DNase I and shaken approximately 30 times to release colonic crypts. The suspension was filtered twice using 100 μm cell strainers. The filtrate was centrifuged for 5 minutes at 1100 RPM, and the pellet was resuspended in 400 to 500 µL of matrigel and plated in 6 independent wells of a 12-well plate. Wild type organoids were cultured in advanced DMEM/F12 (Gibco, 12634028), 10 mM HEPES (Corning, 25-060-Cl), 4 mM L-glutamine (Gibco, 35050061), 0.5 nM WNT Surrogate-Fc fusion protein (ImmunoPrecise Antibodies, N-001), 50 ng/ml EGF (PreProtech AF-100-15-500UG) and RSPO3 conditioned media (2% of final volume). Fungin (InvivoGen, ant-fn-1) and Primocin (InvivoGen, ant-pm-05) were added to the culture for the first seven days.

To generate colon cancer organoids, colon crypts derived from *LSL-KrasG12C* mice were transfected with a construct containing a *Trp53* sgRNA and Cre recombinase, and selected by withdrawing EGF from the media and treating with Nutlin-3 for 10 days^39^. To engineer missense Apc mutations, organoids were nucleofected with a construct containing the base editing enzyme FNLS and a synthetic gRNA targeting codon Q1405, and selected by withdrawing RSPO3 from the media^40,41^. AKP organoids were cultured in murine organoid (MO) medium containing advanced DMEM/F12, 10 mM HEPES, 4 mM L-glutamine, and 50 ng/ml EGF (PreProtech AF-100-15-500UG). Organoid passage and cryopreservation were done as previously described^42,43^. Organoids were passaged in 300 µl of Matrigel plated in one well each of a 6-well dish (Corning 353224). DNA, protein, and RNA isolation were done as previously described^42,43^.

### Patient-derived organoids

Patient-derived organoids were established as previously described44. Patient tumor samples were washed, cut into 5 mm fragments, and incubated in a dissociation buffer containing advanced DMEM/F12 (Gibco, 12634028), 4 mM L-glutamine, 100 U/mL penicillin-streptomycin, 250 U/ml type III collagenase (Worthington, 9001-12-1), and 1 U/ml dispase (Sigma-Aldrich, D4818). The samples were shaken at 37°C for 30 minutes, filtered through a 70 µm cell strainer, centrifuged at 1100 RPM for 5 minutes, and washed with advanced DMEM/F12. Cells were counted and resuspended in Matrigel at a concentration of approximately 10,000 to 50,000 cells per 200 µl of Matrigel in 6-well plates. PDOs were maintained in human organoid (HO) medium, containing advanced DMEM/F12, 10 mM HEPES, 4 mM L-glutamine, 1mM Alonso & Chu et al, Concurrent genetic and non-genetic resistance mechanisms to KRAS inhibition in CRC – preprint N-Acetylcysteine (Sigma-Aldrich A9165), 10 mM nicotinamide (Sigma-Aldrich N0636), 50 ng/ml EGF (PreproTech AF-100-15), 20 ng/mL FGF-10 (PeproTech, 100-26), 10 µM Y-27632 (MedChemExpress, HY-10071), 500 nM A-83-01 (MedChemExpress, HY-10432), 10 µM SB202190 (MedChemExpress, HY-10295), 200 nM LDN193189 (MedChemExpress, HY-12071), 10 nM Gastrin-I (MedChemExpress, HY-P2671), 0.5 nM WNT Surrogate-Fc fusion protein (ImmunoPrecise Antibodies, N-001), RSPO3 conditioned media (2% of final volume), and B27 supplement (Gibco 17504044).

### Organoid growth experiments

Mouse and human colorectal organoids were cultured in Matrigel and maintained in organoid growth media without EGF. Organoids were seeded in 48-well plates. Organoid growth was monitored in real-time using the Incucyte S3 Live-Cell Analysis System (Sartorius). Representative brightfield images were acquired every 24 to 72 hours over a period of 7 days for mouse organoids and 14 days for human organoids. Organoid area was quantified using the organoid module, with automated segmentation and analysis parameters optimized for colorectal organoid morphology. For each time point, organoid area was normalized to the initial area measured on day 0. The following drugs were used: adagrasib, (MedChemExpress HY-130149), RMC-7977 (MedChemExpress HY-148439), TBK1/IKK-IN-5 (MedChemExpress HY-128679), momelotinib (MedChemExpress HY-10961), GSK8612 (MedChemExpress HY-111941), alpelisib (MedChemExpress, HY-15244), defactinib (MedChemExpress HY-12289), TPCA-1 (MedChemExpress HY-10074), pexidartinib (MedChemExpress HY-16749), SP600125 (MedChemExpress HY-12041), LY2157299 (MedChemExpress HY-13226). Drugs were not refreshed during the experiment.

### Patients

All patients received treatment as part of KRAS inhibitor clinical trials approved by the MSKCC Institutional Review Board/Privacy Board under protocols 19-408 (NCT03785249) and 20-183 (NCT04185883). Pathology, radiology reports, medical oncology, and surgery notes were reviewed in the electronic medical record to annotate clinical characteristics for patients. Patient sample collection followed the appropriate Institutional Review Board/Privacy Board protocols and waivers (protocols 06-107, 12-245, 14-019). All participants provided written informed consent for both the clinical trials and biospecimen collection. This study adhered to the ethical principles outlined in the Declaration of Helsinki.

### cfDNA analysis

Circulating free DNA (cfDNA) analysis was conducted using the MSK-ACCESS assay, as previously described16. The MSK-ACCESS assay is a high-depth, custom-designed test that analyzes key exons and domains of 129 genes, along with introns of 10 genes containing recurrent breakpoints. It incorporates duplex unique molecular identifiers and dual index barcodes to reduce sequencing errors and prevent cross-sample contamination. Alterations are identified by comparing against matched normal DNA.

### Patient tissue exome sequencing

MSK-IMPACT, a hybridization capture-based next-generation sequencing assay, was used for sequencing as previously described16,45. Genomic DNA was extracted from formalin-fixed paraffin-embedded (FFPE) tumors and matched normal blood samples using the Qiagen DNeasy Tissue kit and the EZ1 Advanced XL system (Qiagen 69504 and 9018702), respectively. Targeted sequencing was conducted on all exons and selected introns of 468 genes. Sequencing libraries were generated using the KAPA HTP protocol (Kapa Biosystems KK2611) and the Biomek FX system. Captured DNA fragments were pooled into libraries and sequenced on the Illumina HiSeq 2500, achieving high, uniform coverage (>500× median coverage). All genomic alteration types, including substitutions, indels, copy number variations, and rearrangements, were identified by comparing tumor samples to their matched normal counterparts. All sequencing and analyses were performed in a CLIA-certified clinical laboratory.

### Western Blot

Organoids were collected in 5 mL of ice-cold PBS, incubated on ice for 15 min to soften Matrigel, and centrifuged at 2500 RPM for 5 minutes at 4 °C. Organoid pellets were resuspended in 100 µl of RIPA buffer and centrifuged at 500g at 4 °C to collect protein supernatant. Antibodies used for Western blot analysis were: anti-TBK1 (Cell Signaling 3504), anti-Vinculin (Cell Signaling 18799).

### KRAS copy number assay

Organoids were dissociated and pelleted after three days in culture. Following gDNA isolation, copy number assays were performed using the TaqMan Copy Number Assay (Thermo Fisher Scientific, 4400291) according to the manufacturer’s instructions.

### RNA isolation and RNA-seq

Organoids were collected in 5 mL of ice-cold PBS and centrifuged at 2500 RPM for 5 minutes at 4 °C. The pellet was resuspended in 800 µl of TRIzol (Invitrogen, 15596-026). RNA extraction was carried out following the manufacturer’s protocol. To eliminate DNA contamination, the RNA was treated with recombinant DNaseI (Roche Diagnostics, 04716728001) for 15 minutes at room temperature, followed by column purification using the Qiagen RNeasy Mini Kit (74106). cDNA synthesis was performed using 1 μg of RNA and quantified with a NanoDrop spectrophotometer (Thermo Fisher Scientific). The Genomics Core Laboratory at Weill Cornell Medicine assessed RNA quality with an Agilent 2100 Bioanalyzer, prepared the RNA library using the TruSeq Stranded mRNA Sample Library Preparation Kit (Illumina), and conducted RNA sequencing (single-end, 75 cycles) on an Illumina NextSeq 500. Raw FASTQ files have been submitted to the SRA under accession number.

### RNA-seq analysis

Raw FASTQ files were pseudoaligned to the mouse genome (GRCm39) reference using Kallisto (version 0.51.1)^46^. Differential gene expression was estimated using the R package edgeR^47^. Gene set enrichment analysis was performed using the R packages GSEABase and GSVA^48,49^. R (version 4.4.1) and R Studio (version 2024.09.0+375) were used to create all visualizations and principal component analysis. Volcano plots, heatmaps, and other visualizations were produced using ggplot2, Prism (version 10.0.3), and Illustrator (version 28.7.4).

### Spatial transcriptomics

The Integrated Genomics Operation at Memorial Sloan Kettering performed sample processing and data acquisition. Briefly, formalin-fixed, paraffin-embedded (FFPE) tissue sections (5 µm thick) were obtained from 30 patient samples and mounted onto glass slides. FFPE sections underwent deparaffinization and rehydration. The NanoString CosMx platform was used to perform spatial transcriptomics. Tissue sections were subjected to targeted hybridization using the 1000-gene multiplexed human RNA panel. The hybridized samples were then processed following the manufacturer’s protocol.

After hybridization, tissue slides were imaged using the CosMx system, capturing immunofluorescence images and individual RNA transcript location. Antibodies used for immunofluorescence included CD298/B2M membrane marker mix, panCK, CD45, and DAPI (Nanostring). Image processing and transcript localization were performed using NanoString’s pipeline, which includes cell segmentation, transcript quantification, and spatial mapping. Cell segmentation was performed based on nuclear and membrane staining. Raw transcriptomic and spatial data were preprocessed to remove low-quality cells and normalize expression values using the Seurat package50. Cells with less than 100 transcripts were filtered out. Cell clustering and cell typing, gene set enrichment analysis, differential gene expression, spatial transcriptomic images, heatmaps, and volcano plots were generated using Seurat, Prism and Illustrator. Cell-cell interactions were estimated using the package CellChat^31^.

## Statistical analysis

Statistical considerations are reported in each figure legend. Error bars represent s.e.m, unless otherwise noted. In general, student’s t-test (unpaired, two-tailed) was used to assess significance between experimental groups. P < 0.05 was considered statistically significant. Unless otherwise stated, each datapoint represents the average of >3 independent experiments. Experimenters were not blinded to conditions.

## Data availability

Deidentified CosMx data are available at GEO under accession GSE293124. Raw RNAseq fastq files are available at the Sequence Read Archive under accession PRJNA1244364.

---

**Figure S1.**
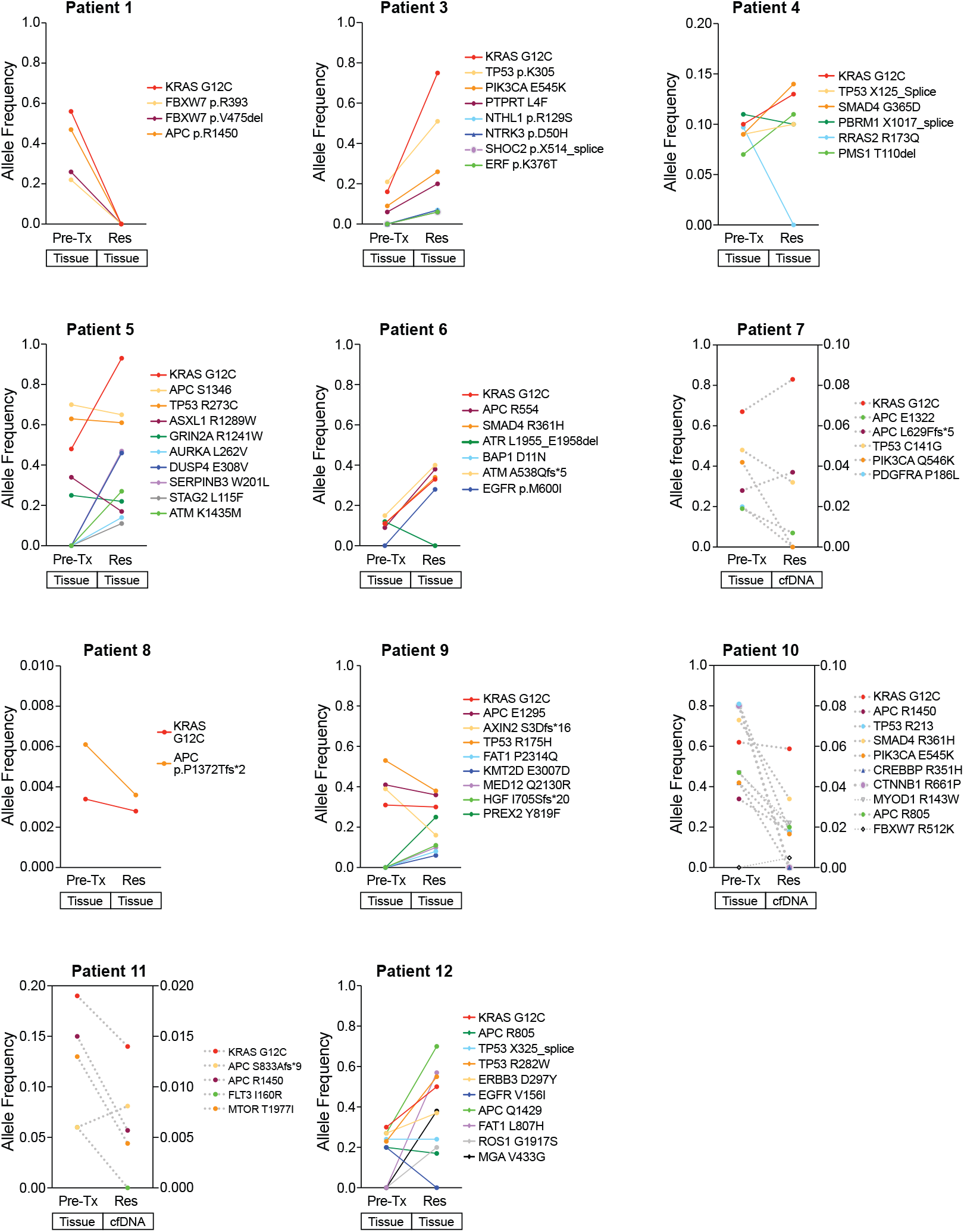
Genomic alterations following treatment with combined KRAS plus EGFR inhibition. Allele frequency of genomic alterations detected in the pre-treatment (Pre-Tx) and resistant (Res) tissue biopsies or cfDNA of patients with CRC treated with KRAS plus EGFR inhibition.

**Figure S2.**
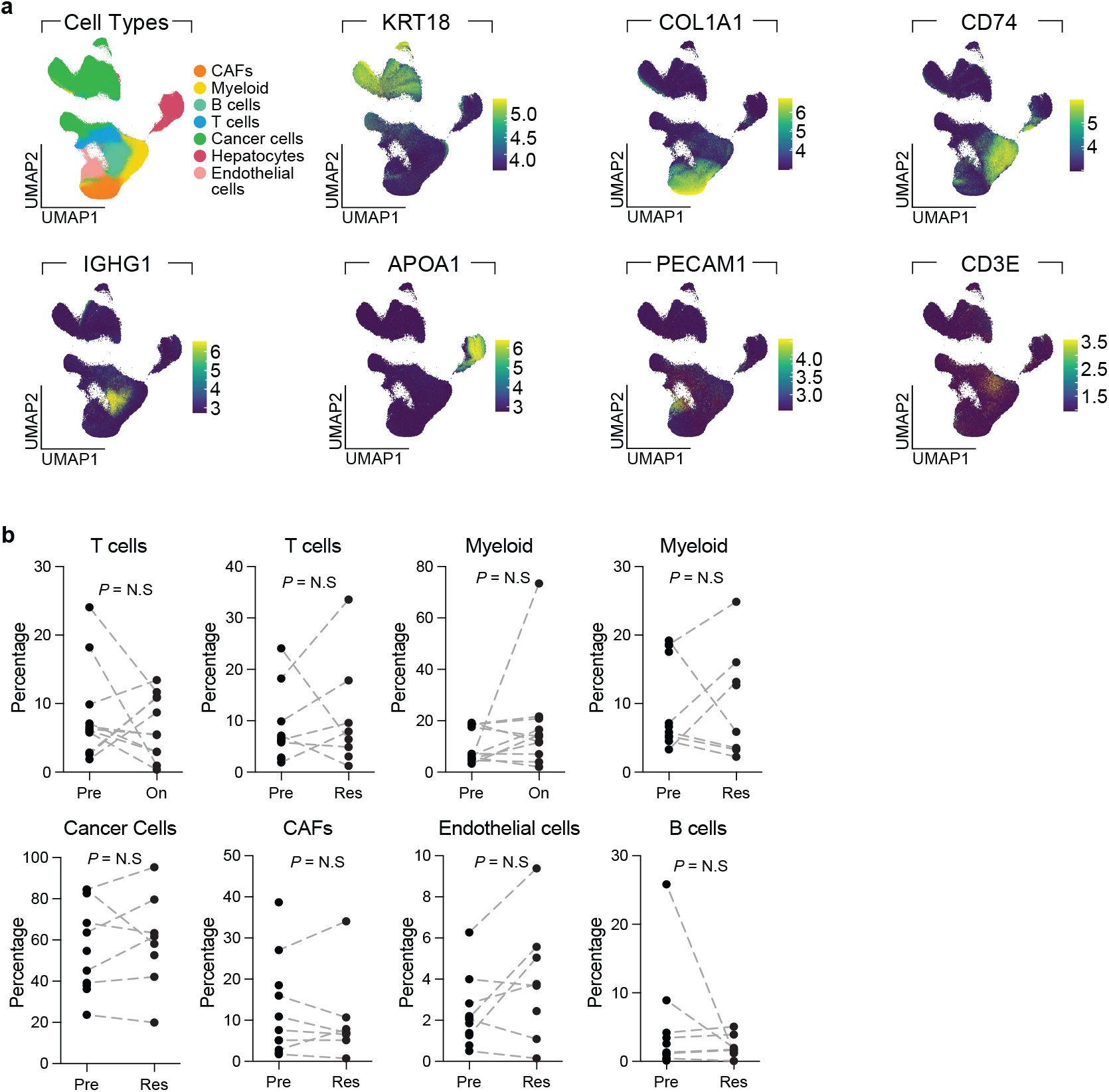
Cellular composition of colorectal tumors treated with KRAS plus EGFR inhibition. **a**, UMAP of patient samples colored by cell type (top left) or gradient showing the log-normalized expression of selected cell typing markers. **b**, Percentage of cell subsets in the pre-treatment (Pre) and on-treatment (On) tissue biopsies. Connected dots represent matched patient samples (N.S. = non-significant).

**Figure S3.**
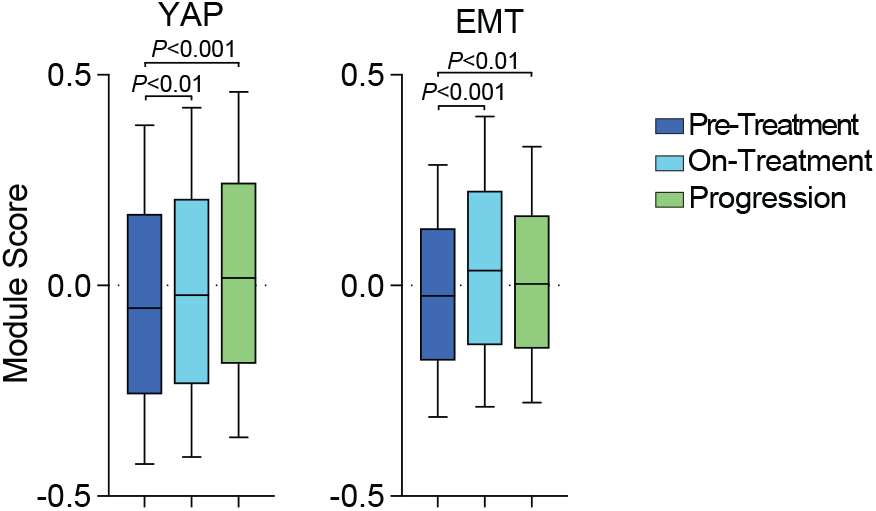
Engagement of resistance adaptive programs by treatment status and treatment response. Module scores of selected gene sets in cancer cells according to treatment timepoint

Error bars show the 10-90 percentiles. *P* values were calculated using a two-tailed Wilcoxon rank sum test.

**Figure S4.**
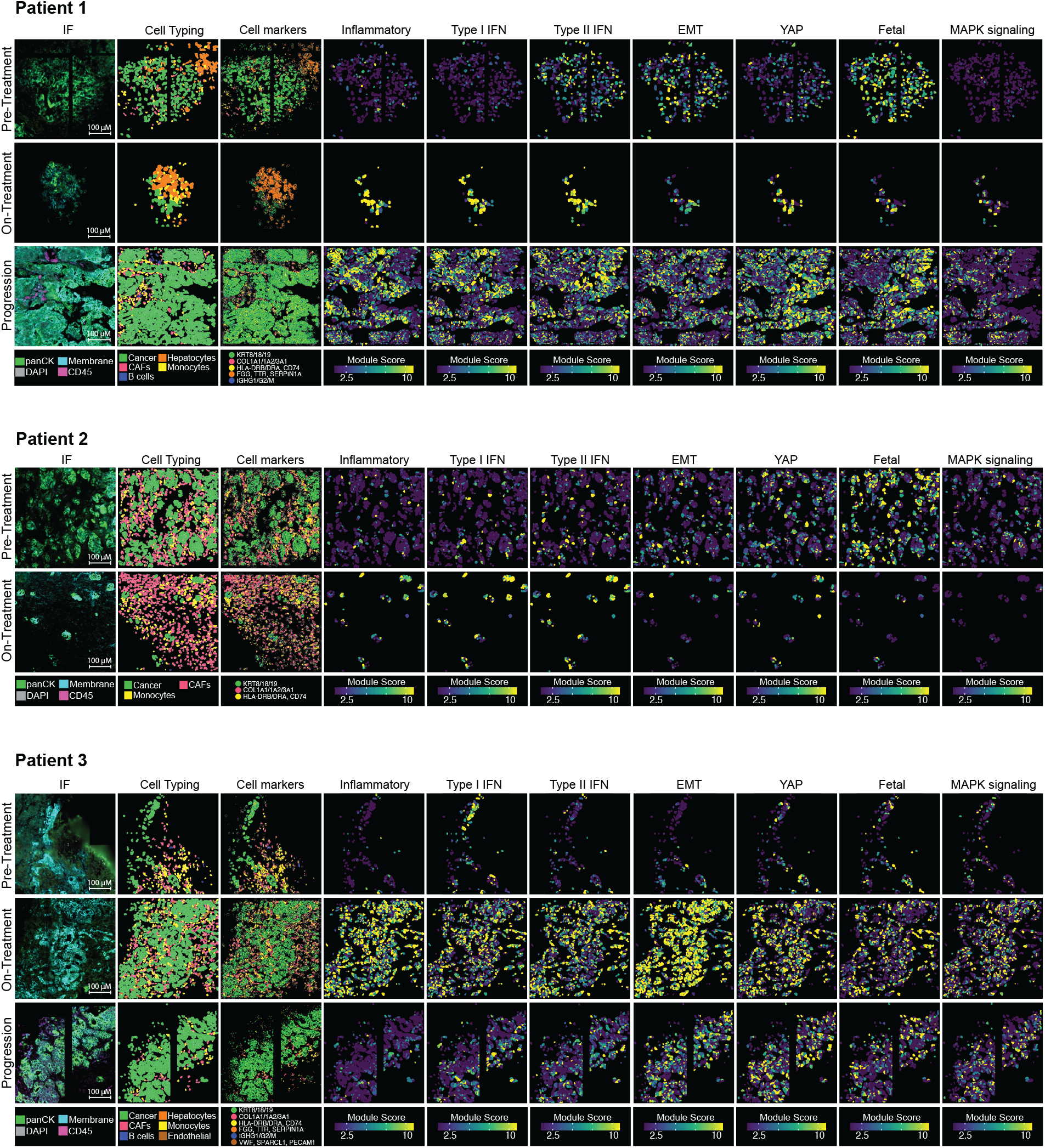
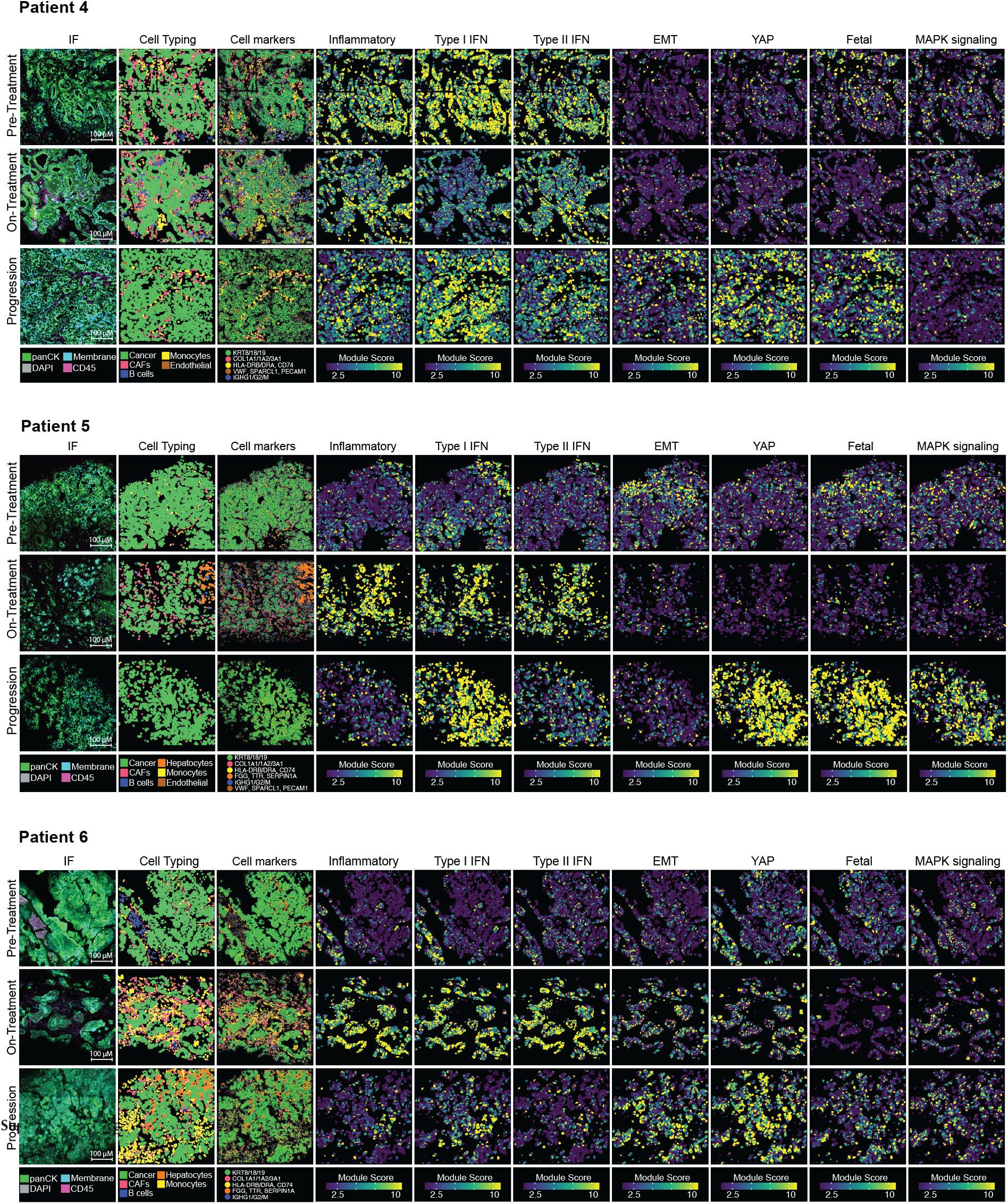
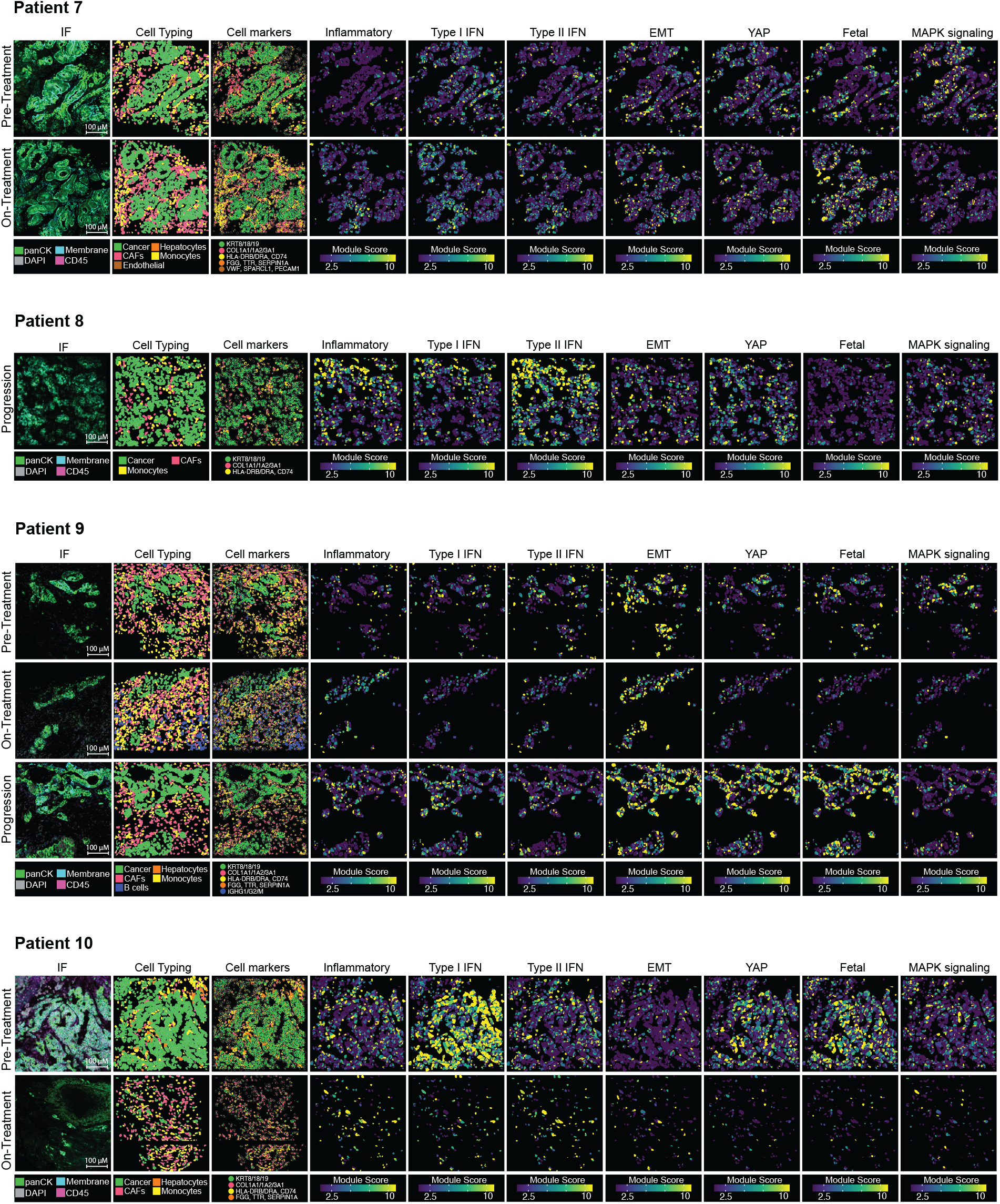
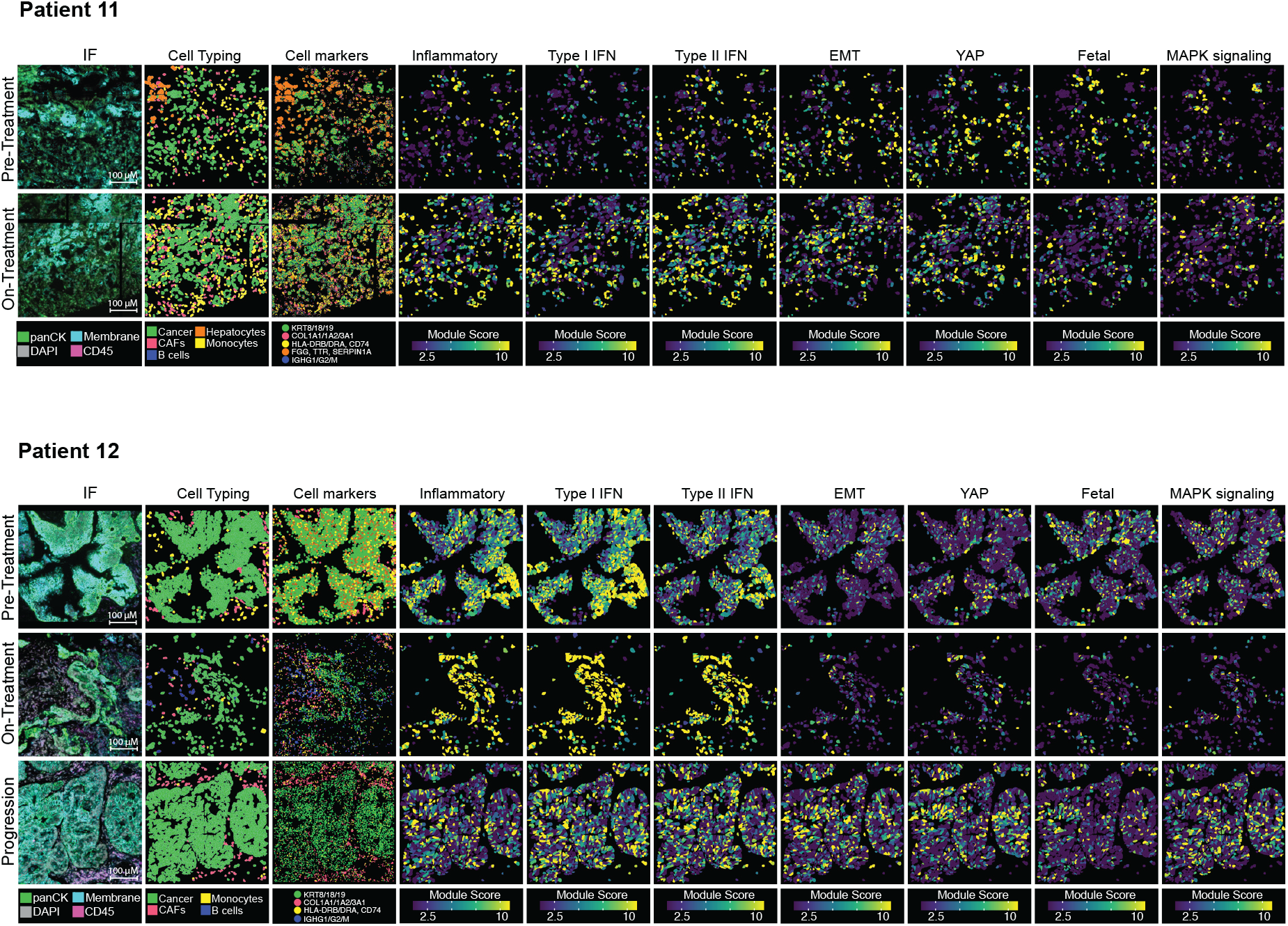
Early and late transcriptional adaptations to KRAS inhibition. Immunofluorescence and spatial images of patient-matched pre-treatment, on-treatment, and post-progression biopsies. The gradient scale shows module score quantiles of resistance programs in cancer cells (non-malignant cells were removed from the analysis).

**Figure S5.**
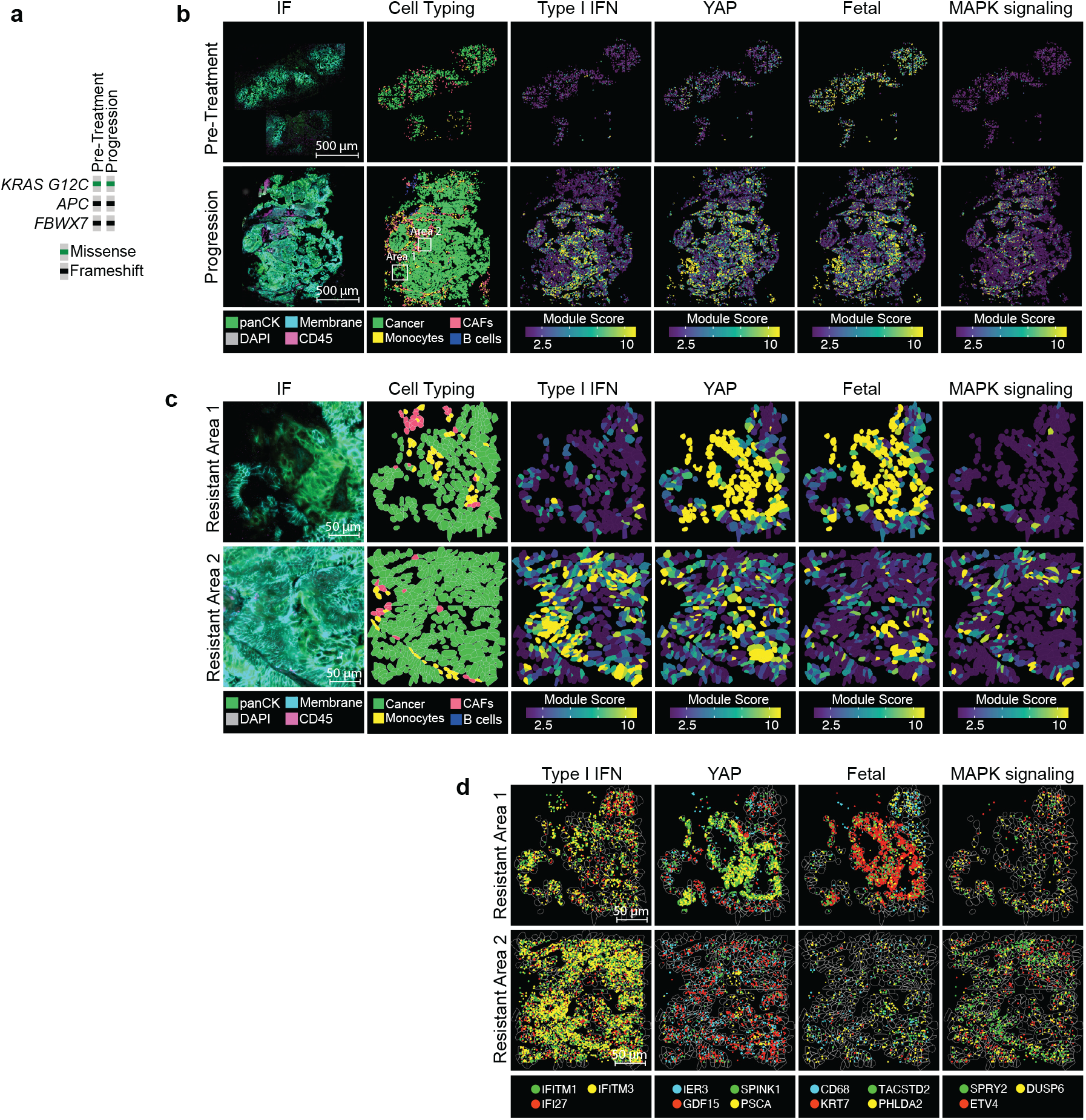
Spatial heterogeneity of adaptive programs. **a**, Oncoprint of the pre-treatment and post-progression biopsies from patient 1. **b**, Immunofluorescence and spatial images of patient 1 observed at low magnification. Two areas in the post-progression biopsy with distinct resistance mechanisms are highlighted. **c**, Area 1 and Area 2 are displayed at higher magnification. The gradient scale shows module score quantiles of selected gene sets in cancer cells. **d**, High-magnification spatial images of Area 1 and 2 showing the location of individual transcripts involved in type I IFN, YAP, fetal and MAPK signaling.

**Figure S6.**
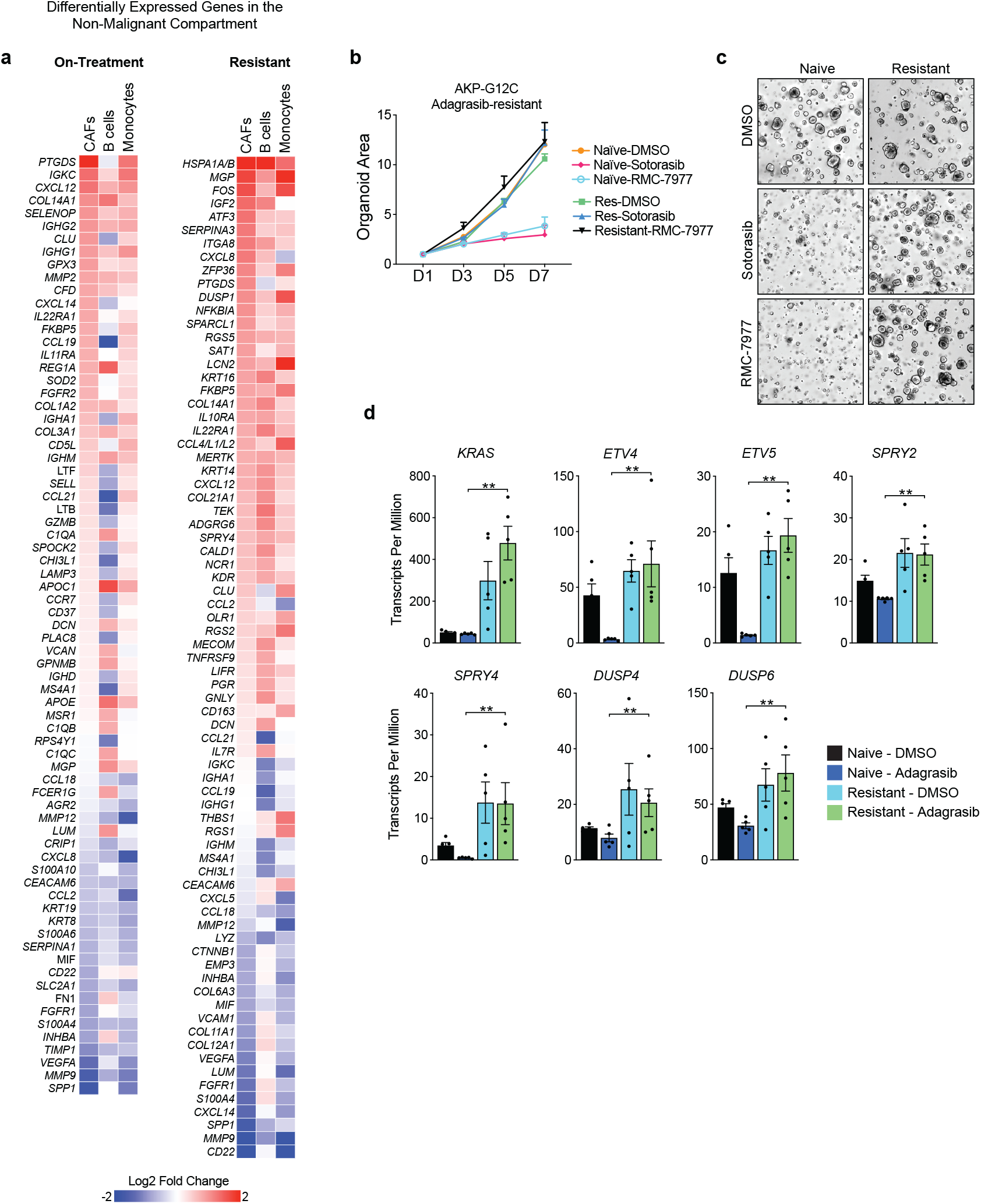
Stromal, immune and cancer cell remodeling following KRAS inhibition. **a**, Differential gene expression in CAFs, B cells, and monocytes in the on-treatment versus pre-treatment, and resistant vs. pre-treatment samples. Genes for which Log2FC > 0.5 or < -0.5 and adj. p < 0.05 are displayed. **b**, Growth of treatment-naïve and resistant AKP-G12C organoids treated with the EC80 of sotorasib (1 µM) or RMC-7977 (20 nM). P values were calculated using a two-tailed t test (n = 4 independent replicates). **c**, Representative brightfield images of treatment-naïve and resistant AKP-G12C organoids treated with the EC80 of sotorasib (1 µM) or RMC-7977 (20 nM). **d**, Normalized expression (transcripts per million) of RAS pathway transcripts in naïve and resistant AKP-G12C organoids treated with DMSO or adagrasib (50 nM for 72 hours). P values were calculated using a two-tailed t test (n = 5 independent resistant lines). ^**^, P < 0.01

**Figure S7.**
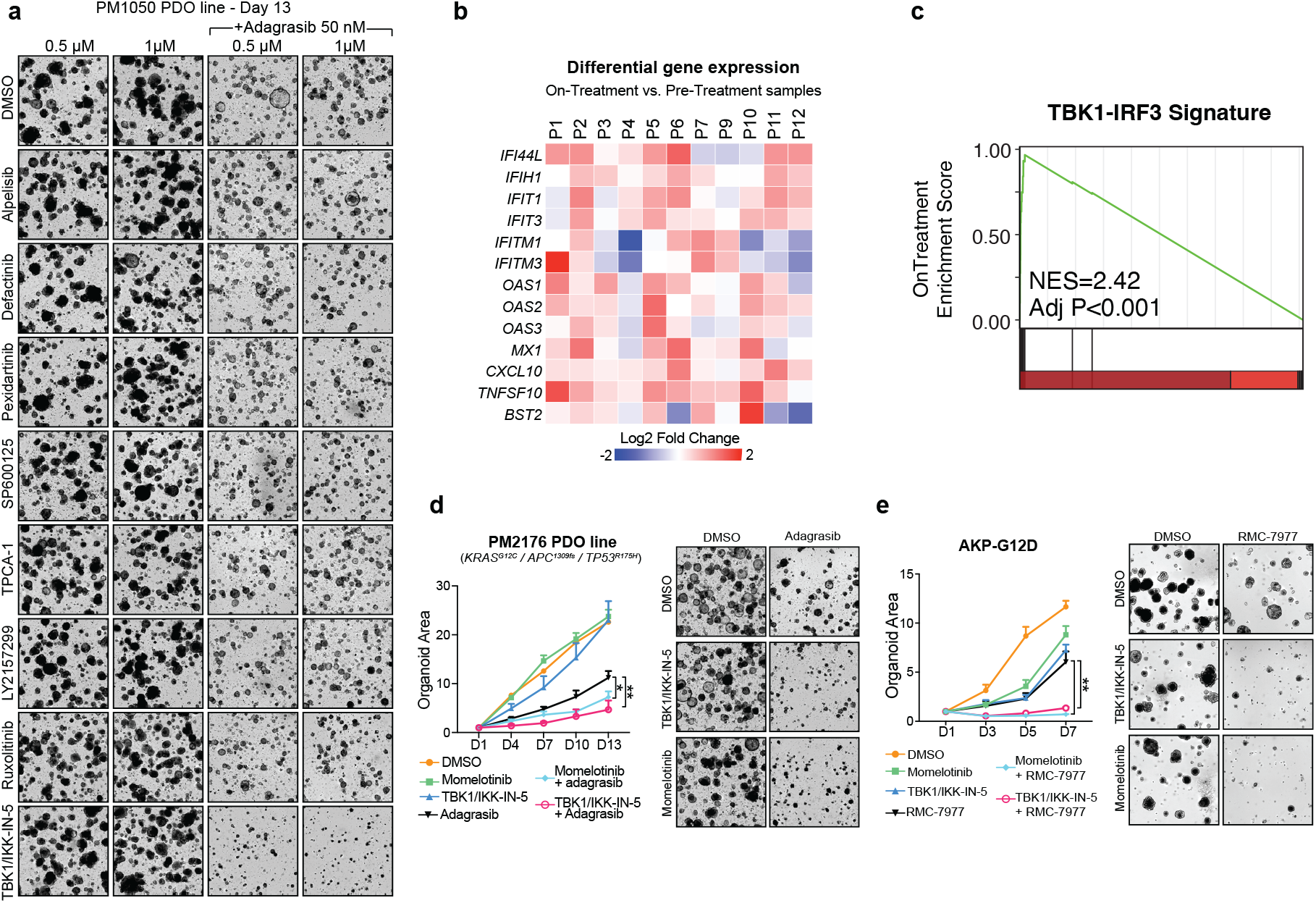
Targeting early inflammatory reprogramming via TBK1 blockage synergizes with KRAS inhibition. **a**, Representative brightfield images of focused drug screen of PM1050 PDOs treated with adagrasib alone (50 nM), kinase inhibitors targeting inflammatory programs (0.5 to 1 µM), or the combination. Two concentrations of each kinase inhibitor were tested. **b**, Cancer cell differential gene expression (Log2 fold-change) of TBK1-IRF3 targets in the on-treatment relative to pre-treatment patient-matched samples. **c**, GSEA plots showing enrichment of TBK1-IRF3 target genes following treatment with adagrasib (50 nM for 72 hours) relative to DMSO-treated AKP-G12C organoids. *P* values were calculated using a permutation test with Benjamini-Hochberg correction. **d**, Growth curve and brightfield images of PM2176 organoids treated with KRASi (adagrasib 50 nM), TBK1i (TBK1/IKK-in-5 1 µM, momelotinib 2 µM), or the combination. **e**, Growth curve and brightfield images of AKP-G12D organoids treated with KRASi (RMC-7977 10 nM), TBK1 inhibition (TBK1/IKK-in-5 2 µM, momelotinib 2 µM), or the combination. *P* values were calculated using a two-tailed t test (n = 4 independent replicates). Error bars represent the standard error of the mean. Brightfield images show organoid density on the final day of the experiment. ^**^, P < 0.01, ^*^, P < 0.05.

## REFERENCES

1. Sung, H., et al. Global Cancer Statistics 2020: GLOBOCAN Estimates of Incidence and Mortality Worldwide for 36 Cancers in 185 Countries. CA Cancer J Clin 71, 209–249 (2021).

2. Siegel, R.L., Giaquinto, A.N. & Jemal, A. Cancer statistics, 2024. CA Cancer J Clin 74, 12–49 (2024).

3. Yaeger, R., et al. Clinical Sequencing Defines the Genomic Landscape of Metastatic Colorectal Cancer. Cancer Cell 33, 125–136 e123 (2018).

4. Canon, J., et al. The clinical KRAS(G12C) inhibitor AMG 510 drives anti-tumour immunity. Nature 575, 217–223 (2019).

5. Hallin, J., et al. The KRAS(G12C) Inhibitor MRTX849 Provides Insight toward Therapeutic Susceptibility of KRAS-Mutant Cancers in Mouse Models and Patients. Cancer Discov 10, 54–71 (2020).

6. Yaeger, R., et al. Adagrasib with or without Cetuximab in Colorectal Cancer with Mutated KRAS G12C. N Engl J Med 388, 44–54 (2023).

7. Fakih, M.G., et al. Sotorasib plus Panitumumab in Refractory Colorectal Cancer with Mutated KRAS G12C. N Engl J Med 389, 2125–2139 (2023).

8. Janne, P.A., et al. Adagrasib in Non-Small-Cell Lung Cancer Harboring a KRAS(G12C) Mutation. N Engl J Med 387, 120–131 (2022).

9. Skoulidis, F., et al. Sotorasib for Lung Cancers with KRAS p.G12C Mutation. N Engl J Med 384, 2371–2381 (2021).

10. Hallin, J., et al. Anti-tumor efficacy of a potent and selective non-covalent KRAS(G12D) inhibitor. Nat Med 28, 2171–2182 (2022).

11. Holderfield, M., et al. Concurrent inhibition of oncogenic and wild-type RAS-GTP for cancer therapy. Nature 629, 919–926 (2024).

12. Kim, D., et al. Pan-KRAS inhibitor disables oncogenic signalling and tumour growth. Nature 619, 160–166 (2023).

13. Popow, J., et al. Targeting cancer with small-molecule pan-KRAS degraders. Science 385, 1338–1347 (2024).

14. Awad, M.M., et al. Acquired Resistance to KRAS(G12C) Inhibition in Cancer. N Engl J Med 384, 2382–2393 (2021).

15. Zhao, Y., et al. Diverse alterations associated with resistance to KRAS(G12C) inhibition. Nature 599, 679–683 (2021).

16. Yaeger, R., et al. Molecular Characterization of Acquired Resistance to KRASG12C-EGFR Inhibition in Colorectal Cancer. Cancer Discov 13, 41–55 (2023).

17. Tanaka, N., et al. Clinical Acquired Resistance to KRAS(G12C) Inhibition through a Novel KRAS Switch-II Pocket Mutation and Polyclonal Alterations Converging on RAS-MAPK Reactivation. Cancer Discov 11, 1913–1922 (2021).

18. Kuboki, Y., et al. Sotorasib with panitumumab in chemotherapy-refractory KRAS(G12C)-mutated colorectal cancer: a phase 1b trial. Nat Med 30, 265–270 (2024).

19. He, S., et al. High-plex imaging of RNA and proteins at subcellular resolution in fixed tissue by spatial molecular imaging. Nat Biotechnol 40, 1794–1806 (2022).

20. Han, T., et al. Lineage Reversion Drives WNT Independence in Intestinal Cancer. Cancer Discov 10, 1590–1609 (2020).

21. Gong, K., et al. EGFR inhibition triggers an adaptive response by co-opting antiviral signaling pathways in lung cancer. Nat Cancer 1, 394–409 (2020).

22. Ruiz-Saenz, A., et al. A reversible SRC-relayed COX2 inflammatory program drives resistance to BRAF and EGFR inhibition in BRAF(V600E) colorectal tumors. Nat Cancer 4, 240–256 (2023).

23. Franca, G.S., et al. Cellular adaptation to cancer therapy along a resistance continuum. Nature 631, 876–883 (2024).

24. Goyal, Y., et al. Diverse clonal fates emerge upon drug treatment of homogeneous cancer cells. Nature 620, 651–659 (2023).

25. Wagner, S., et al. Suppression of interferon gene expression overcomes resistance to MEK inhibition in KRAS-mutant colorectal cancer. Oncogene 38, 1717–1733 (2019).

26. Moorman, A.R., et al. Progressive plasticity during colorectal cancer metastasis. bioRxiv (2023).

27. Mzoughi, S., et al. Oncofetal reprogramming drives phenotypic plasticity in WNT-dependent colorectal cancer. Nat Genet 57, 402–412 (2025).

28. Edwards, A.C., et al. TEAD Inhibition Overcomes YAP1/TAZ-Driven Primary and Acquired Resistance to KRASG12C Inhibitors. Cancer Res 83, 4112–4129 (2023).

29. Chapeau, E.A., et al. Direct and selective pharmacological disruption of the YAP-TEAD interface by IAG933 inhibits Hippo-dependent and RAS-MAPK-altered cancers. Nat Cancer 5, 1102–1120 (2024).

30. Hagenbeek, T.J., et al. An allosteric pan-TEAD inhibitor blocks oncogenic YAP/TAZ signaling and overcomes KRAS G12C inhibitor resistance. Nat Cancer 4, 812–828 (2023).

31. Jin, S., et al. Inference and analysis of cell-cell communication using CellChat. Nat Commun 12, 1088 (2021).

32. Tian, J., et al. Combined PD-1, BRAF and MEK inhibition in BRAF(V600E) colorectal cancer: a phase 2 trial. Nat Med 29, 458–466 (2023).

33. Honda, K. & Taniguchi, T. IRFs: master regulators of signalling by Toll-like receptors and cytosolic pattern-recognition receptors. Nat Rev Immunol 6, 644–658 (2006).

34. Chan, J.M., et al. Lineage plasticity in prostate cancer depends on JAK/STAT inflammatory signaling. Science 377, 1180–1191 (2022).

35. Woolston, A., et al. Genomic and Transcriptomic Determinants of Therapy Resistance and Immune Landscape Evolution during Anti-EGFR Treatment in Colorectal Cancer. Cancer Cell 36, 35–50 e39 (2019).

36. Barbie, D.A., et al. Systematic RNA interference reveals that oncogenic KRAS-driven cancers require TBK1. Nature 462, 108–112 (2009).

37. Zhu, Z., et al. Inhibition of KRAS-driven tumorigenicity by interruption of an autocrine cytokine circuit. Cancer Discov 4, 452–465 (2014).

38. Amen, A.M., et al. Endogenous spacing enables co-processing of microRNAs and efficient combinatorial RNAi. Cell Rep Methods 2, 100239 (2022).

39. Zafra, M.P., et al. An In Vivo Kras Allelic Series Reveals Distinct Phenotypes of Common Oncogenic Variants. Cancer Discov 10, 1654–1671 (2020).

40. Schatoff, E.M., et al. Distinct Colorectal Cancer-Associated APC Mutations Dictate Response to Tankyrase Inhibition. Cancer Discov 9, 1358–1371 (2019).

41. Zafra, M.P., et al. Optimized base editors enable efficient editing in cells, organoids and mice. Nat Biotechnol 36, 888–893 (2018).

42. Dow, L.E., et al. Inducible in vivo genome editing with CRISPR-Cas9. Nat Biotechnol 33, 390–394 (2015).

43. Dow, L.E., et al. Apc Restoration Promotes Cellular Differentiation and Reestablishes Crypt Homeostasis in Colorectal Cancer. Cell 161, 1539–1552 (2015).

44. Chatila, W.K., et al. Integrated clinical and genomic analysis identifies driver events and molecular evolution of colitis-associated cancers. Nat Commun 14, 110 (2023).

45. Cheng, D.T., et al. Memorial Sloan Kettering-Integrated Mutation Profiling of Actionable Cancer Targets (MSK-IMPACT): A Hybridization Capture-Based Next-Generation Sequencing Clinical Assay for Solid Tumor Molecular Oncology. J Mol Diagn 17, 251–264 (2015).

46. Bray, N.L., Pimentel, H., Melsted, P. & Pachter, L. Near-optimal probabilistic RNA-seq quantification. Nat Biotechnol 34, 525–527 (2016).

47. Robinson, M.D., McCarthy, D.J. & Smyth, G.K. edgeR: a Bioconductor package for differential expression analysis of digital gene expression data. Bioinformatics 26, 139–140 (2010).

48. Subramanian, A., et al. Gene set enrichment analysis: a knowledge-based approach for interpreting genome-wide expression profiles. Proc Natl Acad Sci U S A 102, 15545–15550 (2005).

49. Hanzelmann, S., Castelo, R. & Guinney, J. GSVA: gene set variation analysis for microarray and RNA-seq data. BMC Bioinformatics 14, 7 (2013).

50. Satija, R., Farrell, J.A., Gennert, D., Schier, A.F. & Regev, A. Spatial reconstruction of single-cell gene expression data. Nat Biotechnol 33, 495–502 (2015).

